# Genome sequencing of white-blotched river stingray (*Potamotrygon leopoldi*) provides novel clues for niche-adaptation and skeleton formation

**DOI:** 10.1101/2021.10.01.462833

**Authors:** Jingqi Zhou, Ake Liu, Funan He, Yunbin Zhang, Libing Shen, Jun Yu, Xiang Zhang

## Abstract

The white-blotched river stingray *(Potamotrygon leopoldi)* is a cartilaginous fish native to the Xingu River, a tributary of the Amazon River system. It possesses a lot of unique biological features such as disc-like body shape, bizarre color pattern and living in freshwater habitat while most stingrays and their close relatives are sea dwellers. As a member of the *Potamotrygonidae* family, *P. leopoldi* bears evolutionary signification in fish phylogeny, niche adaptation and skeleton formation. In this study, we present its draft genome of 4.11 Gb comprised of 16,227 contigs and 13,238 scaffolds, which has contig N50 of 3,937 kilobases and scaffold N50 of 5,675 kilobases in size. Our analysis shows that *P. leopoldi* is a slow-evolving fish, diverged from elephant shark about 96 million years ago. We find that two gene families related to immune system, immunoglobulin heavy constant delta genes, and T-cell receptor alpha/delta variable genes, stand out expanded in *P. leopoldi* only, suggesting robustness in response to freshwater pathogens in adapting novel environments. We also identified the *Hox* gene clusters in *P. leopoldi* and discovered that seven *Hox* genes shared by five representative fishes are missing in *P. leopoldi.* The RNA-seq data from *P. leopoldi* and other three fish species demonstrate that fishes have a more diversified tissue expression spectrum as compared to the corresponding mammalian data. Our functional studies suggest that the lack of genes encoding vitamin D-binding protein in cartilaginous (both *P. leopoldi* and *Callorhinchus milii)* fishes could partly explain the absence of hard bone in their endoskeleton. Overall, this genome resource provides new insights into the niche-adaptation, body plan and skeleton formation of *P. leopoldi* as well as the genome evolution in cartilaginous fish.

## Introduction

The transition of jawless to jawed vertebrates lays a foundation for the evolution of vertebrates, which was accompanied by many morphological and phenotypic innovations, especially jaws and adaptive immune system [1]. As ancient jawed vertebrates, the fish constitutes a highly diverse and evolutionarily successful class found in both marine and freshwater habitats [2]. The jawed fishes belong to two clades, the cartilaginous fishes (Chondrichthyes) and bony vertebrates (Osteichthyes), which diverged about 450 million years ago (Mya) [3]. Cartilaginous fishes are the most basal group of living fishes, which contain about 1,000 living species [4]. Except for a few published genome information of Chondrichthyes [5–7], too few genetic data in this important taxonomic position were available for scientists to further study the evolution of fishes and the origin of hard bone formation.

The white-blotched river stingray *(Potamotrygon leopoldi),* also known as Xingu River ray, is a freshwater cartilaginous fish native to the Xingu River basin in Brazil [8]. The Xingu River is a geographical part of the Amazon River basin which was inundated by sea during the Pleistocene Epoch. The ancestor of this stingray experienced the transition from marine to freshwater environment. *P. leopoldi* belongs to the family *Potamotrygonidae* in the order *Myliobatiformes* composed of a group of cartilaginous fishes most-closely related to sharks [9]. The species under the family *Potamotrygonidae* all live in the tropical and subtropical regions of South America [9]. Unlike the freshwater stingrays in Africa, Asia and Australia, which belong to the family *Dasyatidae,* most *Potamotrygonidae* species live strictly in freshwater, whereas most *Dasyatidae* species are saltwater dwellers [10, 11]. Except a few widespread members, most river stingrays typically reside in and are confined to a single river basin [10]. For its unique appearance (e.g., white spots on black skin) and distinct behavior (e.g., swimming-maneuvering capabilities), this stingray becomes a pricy pet fish popular in home- and office-based aquaria. Till now, no fish species from the family *Potamotrygonidae* has been extensively studied on genome level. A whole genome data of a *Potamotrygonidae* member and its comparative analysis with other available fish genomes might help us further reveal the evolutionary features unique to *Potamotrygonidae* and provide insights into the ancestral state of gnathostome-specific morphological characters and physiological systems. Therefore, *P. leopoldi* provides an excellent model for studying evolution and niche-adaptation of freshwater cartilaginous fishes.

In this study, we assembled a 4.11 gigabases (Gb) genome of a male stingray, *P. leopoldi,* using the whole genome shotgun (WGS) approach and based on a raw data collection with a total of 370.98× genome coverage, generated from Pacific Biosciences (PacBio) single molecule real time sequencing (SMRT), Illumina HiSeq2000, and 10× Genomics sequencing platforms. We subsequently compared the its genome seqquence to other five representative fish genomes and one chordate genome, including *Cyprinus carpio, Lepisosteus oculatus, Latimeria chalumnae, Danio rerio, Callorhinchus milii, and Branchiostoma Floridae,* to capture its unique evolutionary features and molecular bases. Our results indicate that *P. leopoldi* is one of the slowest-evolving fish species, even within the cartilaginous fish lineage. The transcriptomic data, obtained from six tissues of *P. leopoldi,* shed further lights into highly-diversified gene expression profile among fish lineages, as opposed to the highly-coordinated gene expression among mammalian tissues. The knockout experimentation in fish model reveals the possible genetic foundation for the divergence of hard and cartilaginous skeleton formations. Together, our results start from the *P. leopoldi* genome sequencing to experimental model, providing novel clues for niche adaptation and skeleton formation in the evolutionary history of fish.

## Results

### Sequence and assembly

We constructed a total of 17 sequencing libraries using genomic DNA extracted from a male *P. leopoldi,* and acquired raw data from three sequencing platforms, PacBio SMRT, Illumina HiSeq2000, and 10× Genomics, with coverages of 61.61×, 200.95×, and 108.41 ×, respectively (Table S1). After stringent filtering and redundancy checking, a collective 1,628.64 Gb of sequence data were used for a scaffold-based *de novo* genome assembly. The initial combined assembly was based on the data from long-read PacBio sequences. Paired-end Illumina sequences data and 10× data were used for error correction. The DNA composition of the assembled contigs was of 41.3% GC content (Table S2). The genome sizes estimated by K-mer analysis using Illumina paired-end data were about 4.11 gigabases (Gb) in size with 0.79% heterozygosity (Table S3). The read-to-genome alignment rate is 98.48% with a coverage of 98.74% in the assembly (Table S4). The final assembly consisted of 16,227 contigs and 13,238 scaffolds, a contig N50 of 3,937 kilobases (Kb) and a scaffold N50 of 5,675 Kb in size (Table S5). The completeness of the *P. leopoldi* genome was estimated to achieve 91.8% (3081/3354) coverage using the core benchmarking universal single-copy ortholog (BUSCO) genes. We mapped 248 core eukaryotic genes (CEGs) to the scaffolds and 90% of them are found in predicted exons, based on BLAT scores. Additionally, Merqury gave the accuracy in consensus base calling with 99.9% (Q30) for the genome assembly (Table 1). Thus, BUSCO and CEGs results, and mapping quality indicated that our genome assembly is highly accurate and nearly complete.

**Table 1.** Merqury metrics for drafts of the *P. leopoldi* genome.

| Metric | Genome |
| --- | --- |
| consensus quality value | 37.53 |
| Assembly error rate | 0.01% |
| k-mer completeness (%) | 96.51 |

### Genome annotation

The *P. leopoldi* genome contains more than 71% repetitive content based on *de novo* and sequence homology analyses (Table S6, S7 and Figure S1), which is much higher than that of white shark (58.5%) [12]. A total of 23,240 protein-coding genes were predicted with high-confidence by combining *de novo* and homologous gene prediction methods with the transcriptome data and orthologs from other fish genomes (Table S8). This gene number appears higher than that of elephant shark but lower than that in white shark [1, 12]. We have assigned 23,030 (99.1%) protein-coding genes preliminary functions by referring to BLASTp against protein databases (Swissport, NR, KEGG, and InterPro, Figure S2) and 21,040 (90.5%) predicted genes Gene Ontology (GO) terms, as well as the orthologs to other fish species (Tables S9 and S10). Moreover, noncoding genes including miRNA, tRNA, rRNA, and snRNA, are also identified accordingly (Table S11).

### Phylogenomic and evolutionary analyses

*P. leopoldi* phylogeny is built on a dataset from 24 other fish species and one chordate, i.e., Amphioxus *(B. floridae)* as the outgroup to root our tree, which is composed of 212 one-to-one orthologous genes with 48,202 amino acid sites. A jawless fish, sea lamprey *(Petromyzon marinus),* is basal to cartilaginous and bony fishes and there is a conspicuous split between cartilaginous and bony fishes (Figure 1). Both maximum likelihood and Bayesian trees show exactly the same topology (Figure S3 and S4). *P. leopoldi* is grouped with *C. milii,* forming the Chondrichthyes clade, whereas 22 other bony fishes are clustered as the Osteichthyes clade (Figure S5). This result is consistent with the traditional taxonomic classification of fishes. A MCMCtree-based divergence time estimation indicates that the split of the other fish species from the class Cyclostomata (*P. marinus*) occurred ~533 Mya (Figure 1 and Figure S6), and the splits between Chondrichthyes and Osteichthyes and between the superorder Batoidea (*P. leopoldi*) and Selachimorpha (*C. milii*) are ~381 Mya and ~96 Mya, respectively.

**Figure 1.**
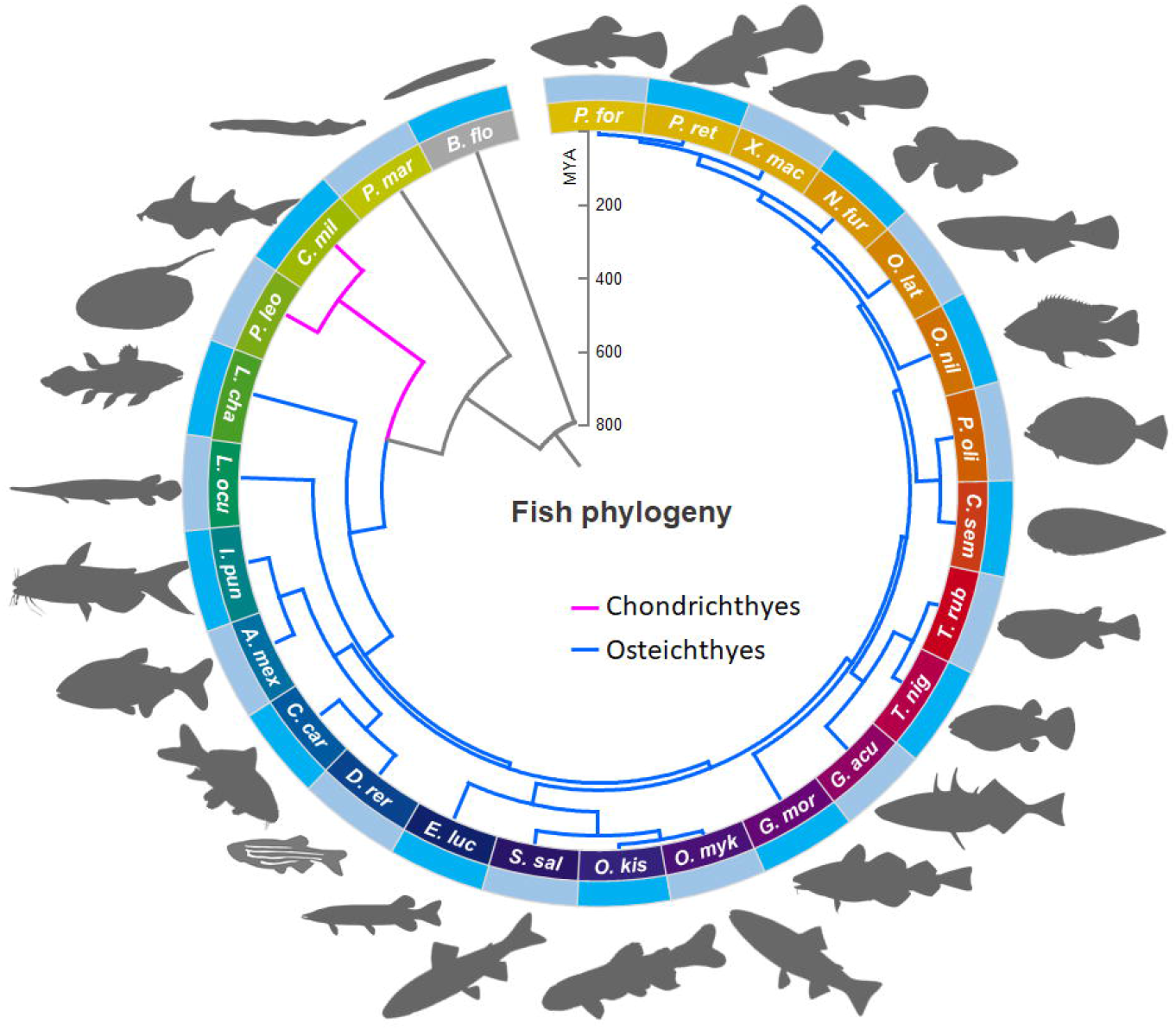
Phylogeny of white-blotched river stingray and 25 selected species. The phylogenomic tree was constructed with FastTree and MrBayes using 212 one-to-one orthologous genes. *B.flo* (amphioxus) was used to root the tree and 25 fishes are *P.mar* (sea lamprey), *C.mil* (elephant shark), *P.leo* (white-blotched river stingray), *L.cha* (coelacanth), *L.ocu* (spotted gar), *I.pun* (channel catfish), *A.mex* (blind cave fish), *C.car* (common carp), *D.rer* (zebrafish), *E.luc* (northern pike), *S.sal* (Atlantic salmon), *O.kis* (coho salmon), *O.myk* (rainbow trout), *G.mor* (Atlantic cod), *G.acu* (three-spined stickleback), *T.nig* (green spotted puffer), *T.rub* (Japanese puffer), *C.sem* (tongue sole), *Poli* (Japanese flounder), *O.nil* (Nile tilapia), *O.lat* (rice fish), *N.fur* (turquoise killifish), *X.mac* (platyfish), *Pret* (guppy), and *P.for* (Amazon molly). Magenta indicates the Chondrichthye lineage and sky blue indicates the Osteichthye lineage. Divergence time is shown in million years (MYA).

We also calculated the evolutionary rates as total substitution rate per site for each species, using the same set of orthologous genes as in our phylogenomic analysis (Table 2). *P. leopoldi* has the lowest amino acid substitution rate among 26 species. Tajima’s relative rate tests confirm that its evolutionary rate is significantly slower than those of *B. floridae, P. marinus, L. chalumnae* and *D. rerio,* but similar to the other cartilaginous fish, such as *C. milii* (Supplemental data 1: Tajima’s rate tests). However, Tajima’s relative rate tests also show that *L. oculatus* exhibits a slower evolutionary rate than *P. leopoldi* when using amphioxus or sea lamprey as the outgroup (Supplemental data 1). Thus, the results above propose that *P. leopoldi* is one of the slowest-evolving fishes.

**Table 2.** Amino acid substitution rate in 26 fish species.

| Species | Number of substitutions per site |
| --- | --- |
| amphioxus | 0.408705 |
| sea lamprey | 0.277635 |
| elephant shark | 0.252635 |
| white-blotched river stingray | 0.250005 |
| coelacanth | 0.265845 |
| spotted gar | 0.250345 |
| channel catfish | 0.338655 |
| blind cave fish | 0.364285 |
| zebrafish | 0.329255 |
| common carp | 0.353285 |
| northern pike | 0.329085 |
| Atlantic salmon | 0.321695 |
| rainbow trout | 0.328565 |
| coho salmon | 0.335825 |
| Atlantic cod | 0.381235 |
| three-spined stickleback | 0.363065 |
| green spotted puffer | 0.410095 |
| Japanese puffer | 0.394355 |
| Nile tilapia | 0.358895 |
| rice fish | 0.407615 |
| turquoise killifish | 0.388785 |
| platyfish | 0.388865 |
| Amazon molly | 0.397335 |
| guppy | 0.393995 |
| tongue sole | 0.391035 |
| Japanese flounder | 0.359955 |

### Orthology and gene family evolution analyses

To explore the evolutionary features of *P. leopoldi,* we performed homology analysis, by comparing its protein coding genes with those of *B. floridae, C. milii, L. chalumnae, L. oculatus, C. carpio* and *D. rerio* (Figure 2a). After filtering the genes shorter than 100 amino acids and with low sequence complexity from the predicted 23,240 protein-coding gene pool, we keep a total of 18,894 stingray protein-coding genes for orthology analysis. Among these kept *P. leopoldi* genes, 12,219, 4,347, and 292 of them are chordate (shared with those of *B. floridae),* bony fish (shared with those of 5 other bony fishes), and cartilaginous orthologs (shared with those of *C. milii),* respectively. In addition, 2,036 of them are *P. leopoldi*-specific (no homologous relationship with the other six species). Notably, *C. carpio* has four rounds of genome duplication and possesses a very large number of protein-coding genes (55,756 protein coding genes) [13].

**Figure 2.**
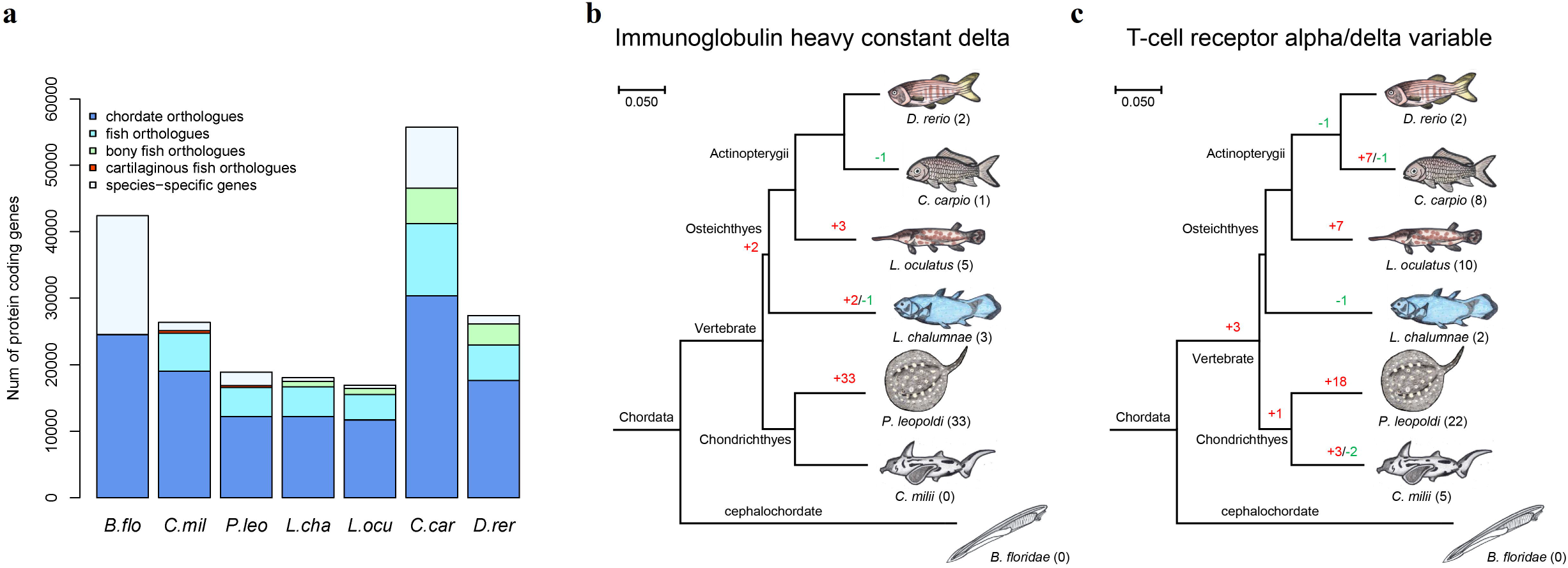
Gene ortholog and family analyses for seven species. **a.** Gene orthology comparison among *B.flo* (amphioxus), *C.mil* (elephant shark), *P.leo* (white-blotched river stingray), *L.cha* (coelacanth), *L.ocu* (spotted gar), *C.car* (common carp), and *D.rer* (zebrafish). **b.** Immunoglobulin heavy constant delta gene expansion and contraction among seven species. **c.** T-cell receptor alpha/delta variable gene expansion and contraction among seven species.

Next, we examined gene family expansions and contractions across seven selected species using a total of 205,728 protein coding genes. According to their homologous relationships, these genes could be classified into 14,818 gene families. 56 gene families are significantly expanded and 14 families are significantly contracted in *P. leopoldi.* GO analyses show that *P. leopoldi* expanded and contracted gene families have different biological emphases (Figure S7). The expanded gene families are primarily related to immune system like defense response to virus (Bonferroni corrected *P* < 0.05), while the contracted gene families are more enriched with cellular component such as crystallin and lectin (Bonferroni corrected *P* < 0.05). Among the *P. leopoldi* gene families experienced expansion, two of them, immunoglobulin heavy constant delta gene family and T-cell receptor alpha/delta variable gene family, are important constituents of vertebrate immune system (Figure 2b and 2c). Interestingly, their expansions are *P. leopoldi*-specific and not found in elephant shark, suggesting their possible roles in freshwater niche adaption. They are also under purifying selection tested with branch model in PAML (data not shown).

### Evolution of *P. leopoldi Hox* gene clusters

The distinct body shape of *P. leopoldi* is assumed to have genetic basis, attributable to its *Hox* gene clusters that exhibit striking spatial collinearity and drive morphologic diversification of almost all metazoans. Due to the contribution of whole genome duplication events among vertebrates (especially in teleost fishes) and lineage-specific secondary losses, the number of *Hox* gene clusters or genes varies greatly among vertebrates [14]. As shown in Figure 3, *Hox* gene clusters range from one in cephalochordate *B. floridae* to 13 in *C. carpio* in number [15, 16], and there are 33 *Hox* genes belonging to four putative *Hox* clusters (A, B, C and D) in *P. leopoldi* within single scaffolds. Therefore, *P. leopoldi* retains the majority 2R *Hox* cluster duplicates. Compared with the *C. milii, P. leopoldi* possesses fewer *Hox* genes, especially in the *HoxC* cluster. Totally, *P. leopoldi* lacks *HoxA4, HoxA5,* and *HoxA6* in the *HoxA* cluster, *HoxB6* in the *HoxB* cluster, *HoxC1, HoxC3, HoxC4, HoxC5, HoxC12* and *HoxC13* in the *HoxC* cluster, and *HoxD4* in the *HoxD* cluster.

**Figure 3.**
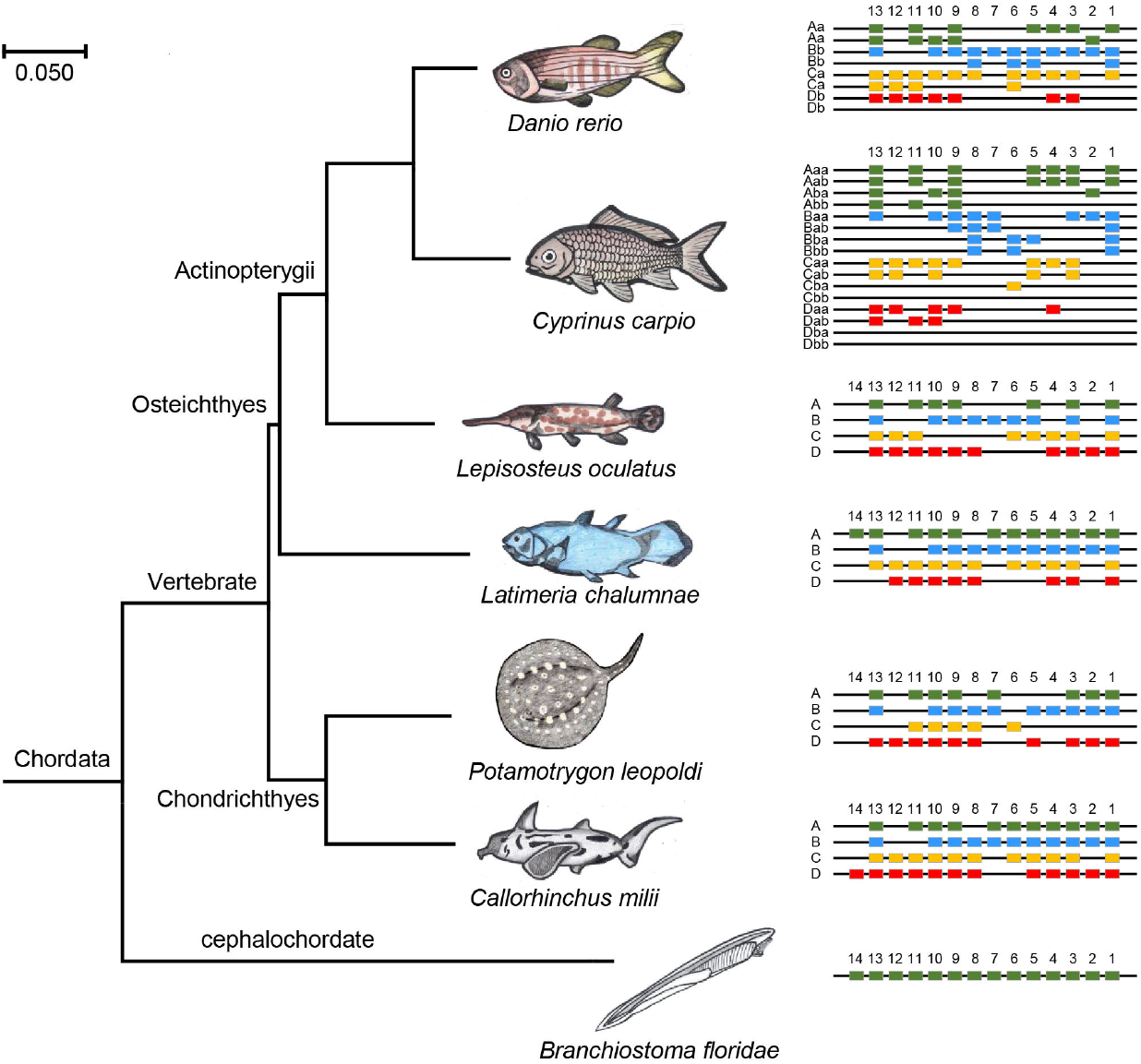
Hox gene clusters identified in *Branchiostoma floridae* (amphioxus), *Callorhinchus milii* (elephant shark), *Potamotrygon leopoldi* (white-blotched river stingray), *Latimeria chalumnae* (coelacanth), *Lepisosteus oculatus* (spotted gar), *Cyprinus carpio* (common carp), and *Danio rerio* (zebrafish).

This difference in *Hox* gene complement may contribute greatly to stingray’s body shape differences compared with elephant shark. Moreover, there have been seven *Hox* genes, including *HoxA5, HoxB6, HoxC3, HoxC4, HoxC5, HoxC13* and *HoxD4,* lost in *P. leopoldi* but present in all other fishes. We assume that such *Hox* gene diversity between *P. leopoldi* and the other fish species may genetically explain its specific body morphology.

### Tissue gene expression profiles

To document its tissue-associated genes and evaluate their possible functions, we acquired RNA sequencing (RNA-seq) data from six *P. leopoldi* tissues. First, we identified each tissue’s differentially expressed genes (DEGs) by comparing its expression profile with the rest five ones. After normalization (transcripts-per-kilobase-million or TPM), 3,559, 4,482, 1,806, 2,347, 1,703, and 2,328 DEGs are found in stingray’s blood, brain, heart, liver, muscle, and skin, respectively. Among these DEGs, 87, 2,280, 184, 537, 388, and 641 genes are up-regulated in stingray’s blood, brain, heart, liver, muscle, and skin, while 3,472, 2,202, 1,622, 1,810, 1,315, and 1,687 are down-regulated in each one of them. Up-regulated DEGs are all tissue-specific and no shared up-regulated genes are found in brain, heart, liver, muscle, and skin (Figure 4a). Gene ontology analyses show that each tissue’s up-regulated DEGs are faithful to their tissue’s function (Table 3). In six tissues, down-regulated DEGs exhibit a similar expression background of antibiotics biosynthesis, catalytic activity and metabolic process (supplemental data). About one third of the up-regulated DEGs in each tissue are *Pl*-species specific (91/218 in blood, 1104/3889 in brain, 361/1109 in heart, 502/1497 in liver, 501/1645 in muscle, and 625/1988 in skin). These *Pl*-specific up-regulated DEGs have no homologous counterpart in *C. milii, L. oculatus,* and *D. rerio* and their possible functions remain to be discovered in stingray.

**Figure 4.**
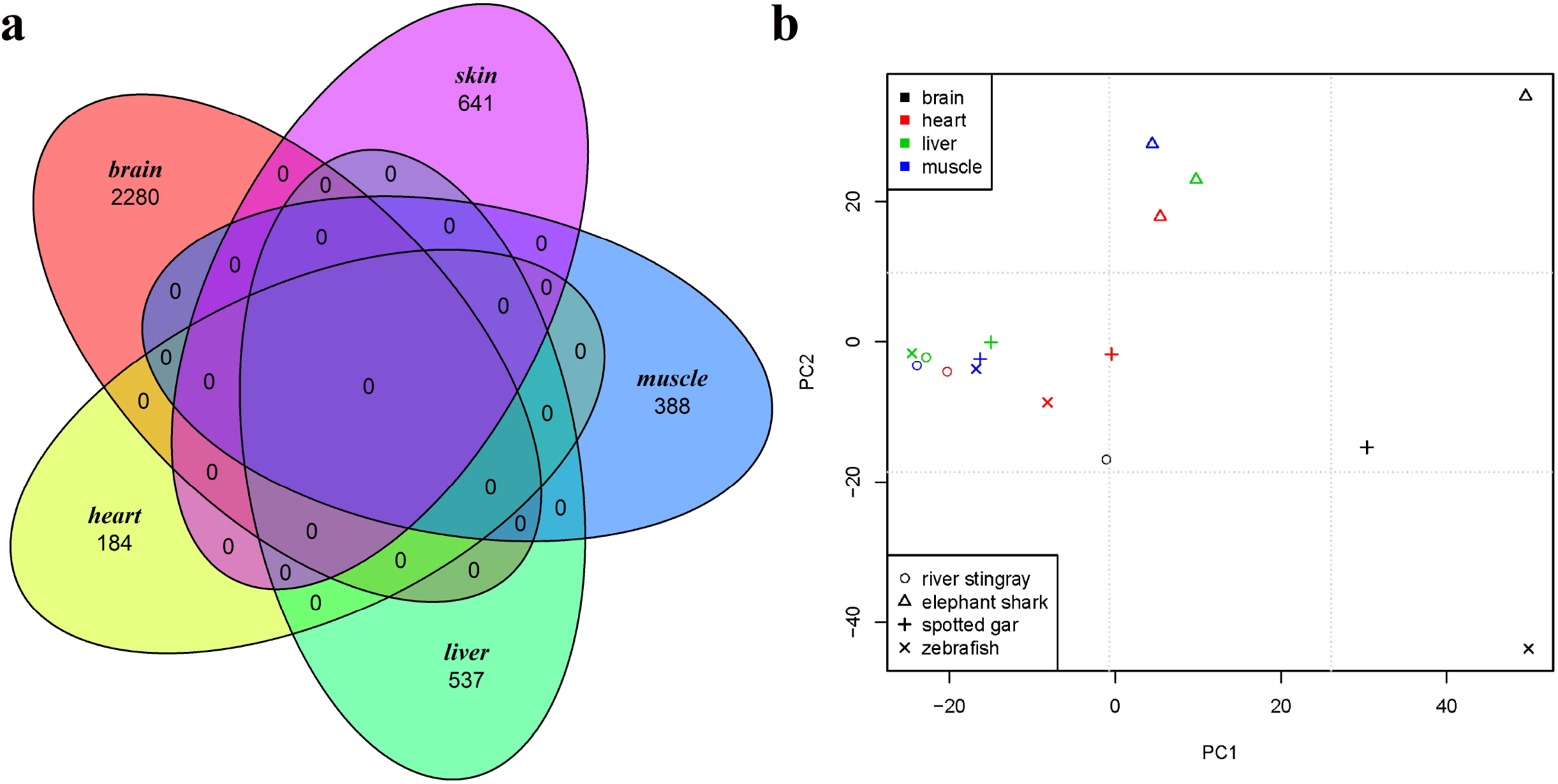
Transcriptome analyses of white-blotched river stingray and other three fishes. **a.** Venn diagram of up-regulated genes in five tissues of white-blotched river stingray. **b.** Principal-component analysis of the expression levels of brain, heart, liver, and muscle using 3738 one-to-one orthologous genes in white-blotched river stingray, elephant shark, spotted gar, and zebrafish.

**Table 3.** DAVID functional annotation analysis of stingray tissue specifically expressed genes.

| Category | Term | P-value |
| --- | --- | --- |
| **Blood** |  |  |
| GOTERM_BP_DIRECT | erythrocyte development | $6.62 \times 10^{-6}$ |
| GOTERM_BP_DIRECT | embryonic hemopoiesis | $3.41 \times 10^{-5}$ |
| GOTERM_BP_DIRECT | erythrocyte differentiation | $6.48 \times 10^{-5}$ |
| GOTERM_BP_DIRECT | hemopoiesis | $7.09 \times 10^{-4}$ |
| **Brain** |  |  |
| GOTERM_BP_DIRECT | central nervous system development | $8.02 \times 10^{-4}$ |
| INTERPRO | Neurotransmitter-gated ion-channel | 0.001797 |
| GOTERM_BP_DIRECT | ganglioside biosynthetic process | 0.002529 |
| GOTERM_BP_DIRECT | neuropeptide signaling pathway | 0.012035 |
| GOTERM_BP_DIRECT | neural crest cell migration | 0.016101 |
| GOTERM_BP_DIRECT | hindbrain development | 0.027042 |
| **Heart** |  |  |
| GOTERM_BP_DIRECT | cardiac muscle cell differentiation | $2.46 \times 10^{-5}$ |
| GOTERM_BP_DIRECT | heart morphogenesis | $6.17 \times 10^{-4}$ |
| GOTERM_BP_DIRECT | heart looping | $9.05 \times 10^{-4}$ |
| GOTERM_BP_DIRECT | heart development | 0.005143 |
| **Liver** |  |  |
| GOTERM_BP_DIRECT | liver development | $7.39 \times 10^{-4}$ |
| GOTERM_MF_DIRECT | oxidoreductase activity | $1.54 \times 10^{-11}$ |
| KEGG_PATHWAY | Glycine, serine and threonine metabolism | 0.007413 |
| KEGG_PATHWAY | Fatty acid degradation | $8.55 \times 10^{-5}$ |
| **Muscle** |  |  |
| GOTERM_BP_DIRECT | muscle organ development | $1.76 \times 10^{-5}$ |
| GOTERM_BP_DIRECT | skeletal muscle tissue development | $7.32 \times 10^{-5}$ |
| GOTERM_BP_DIRECT | myofibril assembly | $2.78 \times 10^{-4}$ |
| INTERPRO | Myogenic basic muscle-specific protein | $4.65 \times 10^{-4}$ |
| GOTERM_BP_DIRECT | skeletal muscle cell differentiation | 0.00223 |
| UP_KEYWORDS | Myogenesis | 0.003216 |
| **Skin** |  |  |
| GOTERM_BP_DIRECT | ectodermal placode formation | 0.068466 |
| GOTERM_CC_DIRECT | melanosome | 0.015904 |

Next, we compared the *P. leopoldi* expression profiles (brain, heart, liver, and muscle) to those of *C. milii, L. oculatus,* and *D. rerio,* using 3,738 one-to-one orthologs among 4 fishes, and performed the principal component analysis (PCA) to investigate the expression patterns for the four tissues across the four fishes. The PCA result shows that a scattered expression pattern for the four tissues across four fishes (Figure 4b). Our data are separated neither by tissue nor by species. In mammals, the expression analyses show that the same tissue from different species tend to cluster together [17], which proposes that the same tissue in different mammals usually perform similar physiological functions. In fishes, the diverged expression pattern of the same tissues suggests a probably much more diversified physiology or a much longer evolutionary history, both of which are not mutually exclusive.

### Genetic basis for hard skeleton formation

As a cartilaginous fish, *P. leopoldi* already has a complex skeleton structure providing support for its body and internal organs.-How hard skeleton emerged in vertebrates and what are the genetic basis for bone formation remain to be investigated. We systematically compared the bone-formation-related gene families between cartilaginous and bony fish genomes. First, we examined the gene family of bone morphogenetic proteins (BMPs) and bone morphogenetic protein receptors among seven selected species. It is known that both BMPs and its receptors play an essential role in skeleton formation. Generally, BMPs are classified into eight different clusters according to their phylogenetic relationship (Figure 5a). Cartilaginous and bony fishes have the representative genes in all 8 clusters. *P. leopoldi* does not have BMP15 gene whereas *C. milii* has. Compared with fishes, *B. floridae* does not have BMP3, BMP9, BMP10, BMP15, and BMP16 genes. As to bone morphogenetic protein receptors [18], both types I and II are present in both cartilaginous and bony fishes but *B. floridae* misses BMP receptor type II (Supplemental data 2: BMPs and BMP receptors in seven species). Our analyses suggest that BMPs and their receptors are less likely to answer the question whether skeleton is made of cartilage or hard bone, since both cartilaginous and bony fishes have all representative BMP and BMP receptor genes. Second, we examined the presence or absence of other bone-formation-related genes between cartilaginous and bony fishes. After scrutinized the candidate gene list involved in skeleton formation from six fish species, we found that vitamin D-binding protein is present in bony fishes but absent in cartilaginous fishes. *L. chalumnae, L. oculatus, C. carpio,* and *D. rerio* have 2, 1, 3, and 1 copies of vitamin D-binding protein, respectively. We hypothesize that vitamin D-binding protein may be involved in the bone formation process for bony fishes. *D. rerio* is a widely used model organism for studies of bone development and formation [19]. To verify the possible function of GC gene (vitamin D-binding protein) in bone formation, we used a sgRNA against EGFP as scrambled control and designed gc-e4 against the exon 4 of GC gene and gc-e8 against exon 8 of GC gene [20]. As shown in Figure 5b and 5c, embryos injected with control RNPs displayed normal bone formation, whereas embryos injected with RNPs against GC gene displayed incomplete craniofacial skeleton mineralization. The results support our hypothesis that GC gene is involved in hard skeleton formation.

**Figure 5.**
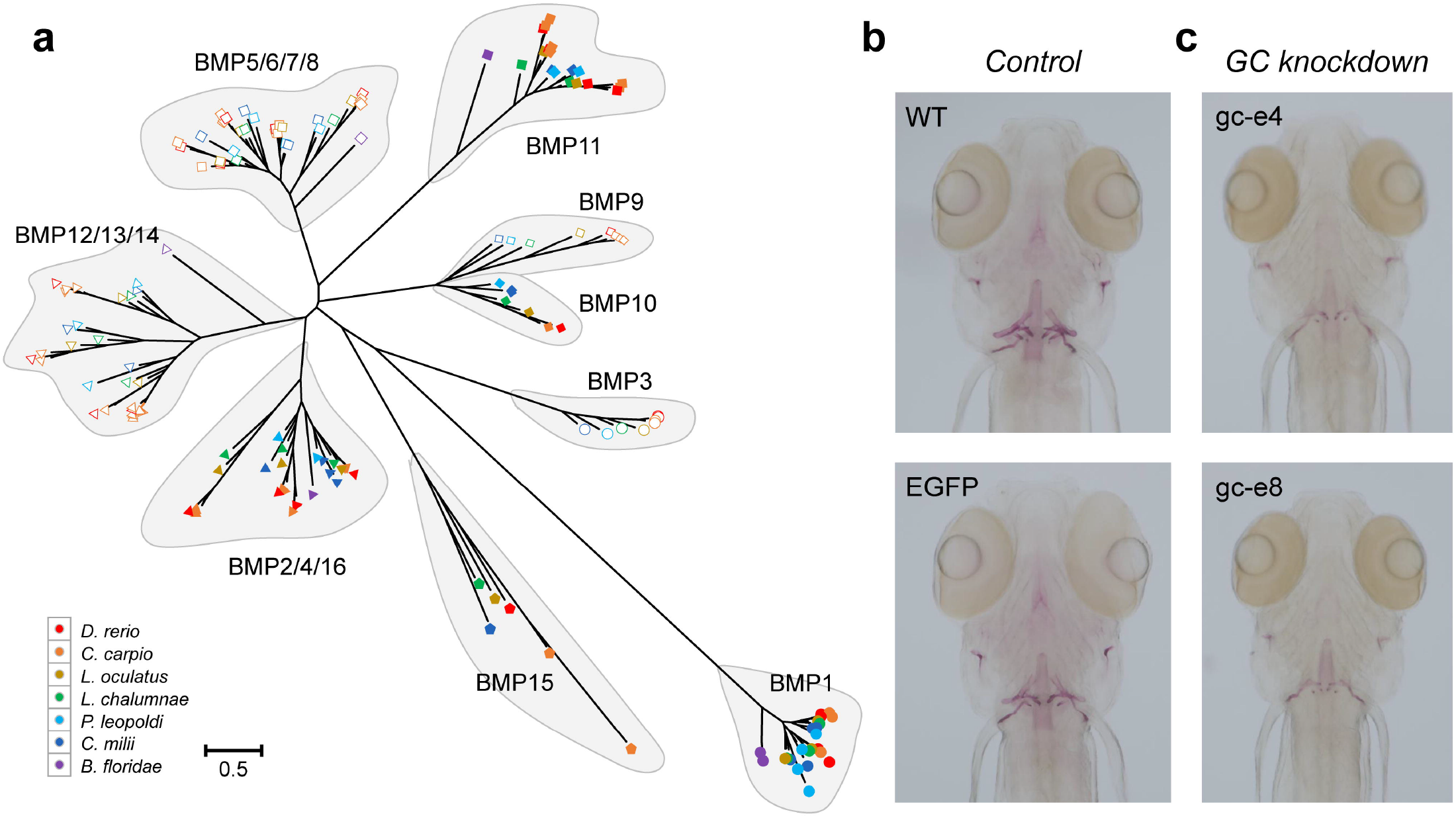
Analyses of bone-formation-related genes among seven selected species. **a.** Phylogenetic analysis of BMP gene family among seven selected species. **b.** Alizarin red staining of 6 dpf control zebrafish (WT: uninjected with RNP, EGFP: injected with enhanced green fluorescent protein RNP). **c.** Alizarin red staining of 6 dpf GC-knockdown zebrafish (gc-e4: injected with GC gene exon 4 RNP, gc-e8: injected with GC gene exon 8 RNP).

## Discussion

### The evolutionary features of *P. leopoldi* genome and Hox gene clusters

The genome data of *P. leopoldi* provide us an information bonanza for understanding fish evolution, especially the split between cartilaginous and bony fishes. Our phylogenomic analyses show that Chondrichthyes and Osteichthyes are two parallel monophyletic groups and the split between them can be dated back to 381 Mya, which is less than the estimation of 450 Mya based on mitochondrial genome [3] The discrepancy between our result and the mitogenomic estimation could be caused by the different evolutionary rates of nuclear genome and mitochondrial genome. Mitochondrion has a particularly high mutation rate and a much diversified evolutionary spectrum across species[21]. The ununified estimations of split time can be explained by different data sources. Generally, the evolutionary rate of elasmobranchs is much lower than that in mammals [22, 23], and the Chondrichthyes also show a slower evolutionary rate than Osteichthyes. In this study, *P. leopoldi, C. milii,* and two basal bony fish species (*L. chalumnae* and *L. oculatus)* evolve much slower than the other fishes in our study (Figure 1). These four fish species also did not experience the third round of genome duplication (3R) happened in the ray-finned fish lineage [24, 25]. This result supports the hypothesis that the 3R might promote the teleost in a higher rate of sequence evolution [1, 26, 27], and suggests that cartilaginous and ancient bony fishes might have been well-adapted to their niches. In *P. leopoldi, C. milii, L. chalumnae* and *L. oculatu,* the substratum (the number of genes) for evolution to work on was limited and therefore slow evolution is expected. Although *P. leopoldi* genome is not assembled at chromosomal level, the completeness of the genome was estimated to achieve 91.8% coverage using BUSCO analysis. *P. leopoldi* has four *Hox* gene clusters containing 33 genes, the smallest number of *Hox* genes among six fishes analyzed in our study (Figure 3). *C. milii, L. chalumnae, L. oculatus, C. carpio,* and *D. rerio* have 45, 43, 34, 62, and 49 Hox genes, respectively. Moreover, *P. leopoldi* lacks *HoxA5, HoxB6, HoxC3, HoxC4, HoxC5, HoxC13* and *HoxD4* presented in the other five fishes. *P. leopoldi* has a disc-like body whereas most fishes usually have a streamline body. The *Hox* gene difference between *P. leopoldi* and the other fishes may help to explain why *P. leopoldi* has a simple body design with large pectoral fins and a whip-like tail. However, our study doesn’t provide the answer for which Hox gene contributes to what morphological alteration in stingray’s body plan.

### The expansion of two immune related gene families *in P. leopoldi*

Strength lies in number. It is also true in gene family evolution. The greater number of genes a family has, the more diversified its function would be. Our gene family analysis shows that 56 gene families were significantly expanded in in freshwater stingray. Many of them are involved in immune and stress response such as heat shock proteins (HSPs), cytochrome P450 (CYP450), inhibitors of apoptosis (IAPs). We found two *P. leopoldi* gene families expanded, directly related to the immune system, which are immunoglobulin heavy constant delta (IGHD) and T-cell receptor alpha/delta (TCR α/δ) variable (V) genes (Figure 2). Immunoglobulins are membrane-bound or secreted glycoproteins produced by B lymphocytes. Immunoglobulin D (IgD) is made up of two heavy chains of the delta class, and two light chains. IgD is present in species ranging from fish to mammal, suggests that IgD has important immunological functions. IgD has been reported be able to bind to basophils and mast cells and induce antimicrobial factors, which contributes to immune surveillance and inflammation under pathological conditions [28]. TCR α recognize the peptides that are bound to major histocompatibility complex (MHC) molecules and TCR δ recognize antigens directly, which are considered as a bridge between the innate and adaptive immune system. TCRα and TCRδ cDNA sequences have been identified in the nurse shark [29], as well as in other vertebrates. The existence of TCRs in sharks suggests that the adaptive immune system evolved in cartilaginous fish with the same fundamental major components that exist in vertebrates such as humans [30]. Our studies of TCR expansion in *P. leopoldi* help elucidate that the V genes of TCRs have evolved for 500 million years, and indicate their role in the diversification of the pre-immune repertoire. The natural habitat of *P. leopoldi* is Xingu River basin and the phylogenetic relationship of stingrays shows that the family *Potamotrygonidae* evolved from the sea-dwelling ancestors [31]. The fact that stingrays can transit from marine environment to pathogen rich freshwater suggests stingrays should have a complex immune system. The expansion of immune system-related genes (Figure 2) suggests that *P. leopoldi* could require specific immune responses for pathogens and antigens in new niches, although there has been no evidence that these two gene families are directly involved in freshwater habitat adaptation. Selective pressure analyses further show that they are under purifying selection, a sign of function constraint and importance. It indicates that *P. leopoldi* fast adapted to its new niche in Xingu River after its ancestor migrated from the Atlantic Ocean to the Amazon River system.

### The gene expression silhouette of *P. leopoldi* tissues

Expression profile comparison among five *P. leopoldi* tissues demonstrates that its brain has a much wider expression spectrum than the other tissues (Figure 4), and the result is consistent with that of mammalian organs. In mammals, central nervous system has more specifically expressed genes than heart and liver [32]. Compared with mammals, there are more species-specific genes expressed in *P. leopoldi* in each of the analyzed tissues [17]. The brain-specific modules are enriched with genes involved in typical processes as for central nervous system development (Benjamini–Hochberg corrected *P*□ <□0.05), and thus define common neural tissue functions. Furthermore, the same tissue from different fishes exhibits a much more diversified expression pattern than its counterpart in mammals. The earliest mammal appeared around 225 Mya and our analysis shows that fishes emerged at least 533 Mya [33]. As fishes have a much longer evolutionary history than mammals, a greater divergence for fishes on both expression and sequence levels is expected. We are only able to identify 3,738 one-to-one orthologous genes among stingray, elephant shark, coelacanth and zebrafish, which is much less than 5,636 amniotic one-to-one orthologous genes identified among nine endothermic species in a previous study [17]. Therefore, the long fish evolutionary history may explain the non-unified expression profiles of their tissues at least in part. Our analysis of tissue transcriptomes from all representative fish lineages refines previous hypotheses and provides a new viewpoint for the evolution of chordate tissue functions.

### Absence of vitamin D binding protein in *P. leopoldi*

Skeleton is an essential part of vertebrate body, which can be made of hard bone or cartilage only. For bone, two cell types, osteoblast and osteocyte, contribute to the formation and mineralization, but cartilage has only one cell type called chondrocyte [18, 19]. Therefore, the emergence of hard skeleton in vertebrates must have engaged complex cellular processes and multiple genetic inventions. We searched the *P. leopoldi* gene inventory for the ones known to be involved in bone formation in osteichthyans. All gene families seem to involve in skeleton formation were present, except the vitamin D-binding protein (VDP). Our analyses of BMPs and BMP receptors show that both cartilaginous and bony fishes have all representative members of these two gene families (Figure 5a). The slight difference of BMPs and BMP receptors between the two fish lineages is insufficient to explain the major departures in their skeletons. One major difference between bones and cartilages is whether calcium phosphate is present in their extracellular matrix or not. The previous study has shown that secreted calcium-binding phosphoproteins involve in the bone formation in *D. rerio* [1]. The analysis of *C. milii* genome has reported the lack of genes encoding secreted calcium-binding phosphoproteins (SCPP) in cartilaginous fishes, which explained the absence of bone in their endoskeleton [1]. Our study further demonstrates that vitamin D-binding protein (DBP) may also take part in bone formation process (Figure 5b, c). Vitamin D promotes the absorption of calcium through intestines [34]. Together, our result and that of a previous study both suggest that genes responsible for calcium metabolism is essential for the hard-skeleton formation in bony fishes. Since cartilaginous and bony fishes evolved in parallel, several crucial genes alone may sufficiently exclude the hard skeleton from cartilaginous fishes, albeit under ongoing studies.

## Conclusion

In this study, we report the generation and analysis of a draft genome sequence of *P. leopoldi,* a cartilaginous freshwater fish. Because cartilaginous fishes constitute a critical outgroup for understanding the evolution and diversity of bony vertebrates, the whole-genome analysis shows that the *P. leopoldi* genome is evolving significantly slower than other vertebrates. The transcriptomic data shed lights into highly-diversified gene expression profile among fish lineages, as opposed to the coordinated gene expression among mammalian tissues. The expansion of immune related genes IGHD and TCRs suggests that the diversification of the pre-immune repertoire in cartilaginous fish could play a role in the evolution of an adaptive immune system. Our study further demonstrates that vitamin D-binding protein (DBP) may partly explained the absence of hard bone in their endoskeleton. Together, our results starting from the *P. leopoldi* genome to experimental model provide novel clues for niche adaptation and skeleton formation in the evolutionary history of fishes.

## Materials and Methods

### *P. leopoldi* sample

A mature male *P. leopoldi* individual was acquired from an aquarium in China in January, 2018. It was the descendant of captive *P. leopoldi* breeding population. The fish was killed in a humane way and the experimental procedure was performed in accordance with the guidelines of the Animal Care Committee at the Institute of Neuroscience, Shanghai Institutes for Biological Sciences, Chinese Academy of Sciences. Its skin, heart, blood, muscle, liver, and brain were used for DNA and RNA preparation and sequencing libraries construction.

### Genome sequencing and assembly

The genomic DNA of *P. leopoldi* was sequenced by whole genome shotgun strategy. Based on the genome features, three different lengths (230bp, 350bp, and 450bp) of DNA inserts were produced. The Illumina HiSeq2000 platform was used to sequence these reads by the pair-end sequencing method with the read length of 150 bp in order to capture the whole genome data. A total of 17 DNA libraries were constructed and the total amount of sequencing data were 882.2 Gb with a coverage of 200.95×. The PacBio SMRT platform (yielding an average read length of 20 Kb) was also used to generate 270.51 Gb data, equivalent to a genome coverage of 61.61×. Additionally, a 10× Genomics library was constructed, coupled with the Illumina sequencing platform in a read length of 150 bp, yielding 475.93 Gb data equivalent to a genome coverage of 108.41 ×. PacBio long reads were utilized to perform *de novo* assembly. Around 31 million subreads were used for the assembly with FALCON v0.3.0 to generate contigs [35]. Primary contigs were polished using Quiver5. The scaffolds were built based on 10× Genomics data. Sequence data were generated using the 10XG GemCode platform and the error corrected contigs were used as input for scaffolding to obtain the primary assembly. After scaffolding, shotgun sequences were used to close gaps between contigs. Paired-end clean reads from the Illumina platform were aligned to the assembly with BWA [36]. Base-pair correction was performed for the assembly by using Pilon24. Pilon mostly corrected single insertions and deletions in regions enriched with homopolymer. Contigs or scaffolds shorter than 10 kb were excluded from the analysis to avoid spurious misassembly. Gaps in contigs and scaffolds were closed with subreads. To survey the characteristics of the genome, a total of ~140 Gb next-generation sequencing data equivalent to a genome coverage of 33× were generated. Adaptor sequences, PCR duplicates, and low-quality sequences were removed from the raw data in order to generate high-quality sequences. K-mer statistics of the high-quality sequences were calculated by Jellyfish (version 2.2.7) with “-G 2 -m 17” parameters [37]. GenomeScope 2.0 (https://github.com/tbenavi1/genomescope2.0) [38] was used to estimate the size, heterozygosity rate, and repeat content of the *P. leopoldi* genome. Finally, the completeness of the assembly was assessed through BUSCO analysis (Version 5.2.1, https://busco.ezlab.org/) and core eukaryotic genes (CEGs) analysis (http://korflab.ucdavis.edu/Datasets/cegma/).

### Genome annotation

Repeat elements were annotated with both homology annotation and *de novo* prediction. RepeatMasker and RepeatProteinMask (http://www.repeatmasker.org/RepeatProtein-Mask.html) were used to search the assembled genome against RepBase for known repeat elements [39, 40]. LTR-FINDER and RepeatModeler (http://www.repeatmasker.org/RepeatModeler.html) were used to *de novo* develop repeat element library [41]. After the library was established, RepeatMasker was further used to detect species-specific repeat elements. Tandem repeats were also searched with TRF in the assembled *P. leopoldi* genome [42]. Overlapping transposable elements belonging to the same type of repeats were integrated together.

Protein-coding genes were predicted through combination of *de novo* annotation, homology annotation, and transcriptome-based annotation. Augustus, GlimmerHMM, SNAP (http://homepage.mac.com/iankorf/), Geneid, and Genscan (https://www.genes.mit.edu/GENSCAN.html) software packages were used to *de novo* predict protein coding genes in the *P. leopoldi* genome [43–45]. For homology annotation, the protein sequences from Japanese puffer *(Takifugu rubripes*) rice fish *(Oryzias latipes),* Nile tilapia *(Oreochromis niloticus),* Atlantic cod *(Gadus morhua),* elephant shark (Callorhinchus milii), green spotted puffer *(Tetraodon nigroviridis),* zebrafish *(Danio rerio),* amphioxus *(Branchiostoma Floridae),* three-spined stickleback *(Gasterosteus aculeatus),* and coelacanth (Latimeria chalumnae) were used to search the homologous genes in *P. leopoldi* genome with BLAST and GeneWise softwares [46, 47]. For transcriptome based annotation, the RNA-seq reads from *P. leopoldi* skin, heart, blood, muscle, liver, and brain were mapped and assembled with PASA and cufflinks [48, 49]. EVidenceModeler was employed to integrate the gene sets from three annotation methods into a complete and non-redundant gene set [50]. Finally, PASA was used to correct the EVM annotation result with UTR and alternative splicing information.

Function annotation was performed through comparing the annotated protein coding genes with the known protein banks. The final gene set was blasted against four common protein banks, SwissProt (http://www.uniprot.org/), NR (http://www.ncbi.nlm.nih.gov/protein), KEGG (http://www.genome.jp/kegg/), and InterPro (https://www.ebi.ac.uk/interpro/). IPRSCAN was used to integrate the functional results from four protein banks [51].

The tRNA genes were identified by tRNAscan-SE software with eukaryote parameters [52]. The rRNA fragments were predicted by aligning to whale shark and *C. milii* template rRNA sequences using BlastN at E-value of 1e-10 [53]. The miRNA and snRNA genes were predicted using INFERNAL software by searching against the Rfam database (release 9.1) [54].

### Ortholog analysis

We first compiled the complete proteomes of 25 fish and one chordate genomes. Proteomes for 13 selected organisms included blind cave fish *(Astyanax mexicanus),* zebrafish *(Danio rerio),* Atlantic cod *(Gadus morhua),* three-spined stickleback *(Gasterosteus aculeatus),* coelacanth *(Latimeria chalumnae),* spotted gar *(Lepisosteus oculatus),* Nile tilapia *(Oreochromis niloticus),* rice fish *(Oryzias latipes),* sea lamprey *(Petromyzon marinus),* Amazon molly *(Poecilia formosa),* Japanese puffer *(Takifugu rubripes),* green spotted puffer *(Tetraodon nigroviridis),* and platyfish *(Xiphophorus maculatus)* were obtained from the Ensembl database (release 83, https://www.ensembl.org/). Another 11 proteomes including elephant shark *(Callorhinchus milii),* tongue sole *(Cynoglossus semilaevis),* common carp *(Cyprinus carpio),* northern pike *(Esox Lucius),* channel catfish *(Ictalurus punctatus),* turquoise killifish *(Nothobranchius furzeri),* coho salmon *(Oncorhynchus kisutch),* rainbow trout *(Oncorhynchus mykiss),* Japanese flounder *(Paralichthys olivaceus),* guppy *(Poecilia reticulate),* and Atlantic salmon *(Salmo salar)* were downloaded from the NCBI database (https://www.ncbi.nlm.nih.gov/). The proteome of *B. floridae* was download from JGI Genome Portal (https://genome.jgi.doe.gov/portal/). Proteins shorter than 100 amino acids were discarded and, for alternatively spliced genes, only the longest splice variant of each gene was retained.

Orthologous protein groups were determined with Orthofinder V.2.3.3 with blast search and default parameters [55]. This procedure led to 2,792 orthologous groups with at least one representative gene from 25 species above plus *P. leopoldi*. To maximize orthology, these orthologous groups were filtered with an in-house Perl script to provide a subset group that contained strict one-to-one orthologous gene from each species. We eventually retained 212 one-to-one orthologous groups for phylogenomic analysis.

### Phylogenomic analysis and estimation of divergence time

Protein sequences in each of the 212 one-to-one orthologous groups were aligned using Muscle and ClustalW and the resulting alignments were combined by M-Coffee to produce the multiple sequence alignments (MSA) [56–58]. We used an in-house Perl script to remove gaps and the final MSA contained 48,202 amino acid sites. FastTree 2.1 was used to construct the maximum likelihood tree for the final MSA with JTT and CAT model [59]. MrBayes 3.2.6 was used to construct the Bayesian tree for the final MSA with JTT and invgamma model [60]. We ran MCMC algorithm for 500,000 generations with 4 chains. Bayesian trees were sampled every 100 generations and the first 25% of trees were excluded from the analysis as burn-in. The Bayesian tree was summarized after the average standard deviation of split frequencies below 0.01. RAxML was used to constructed a maximum likelihood using JTT model with gamma distribution [61]. Then, MCMCtree was used to predict the divergence time of 26 species [62]. The intervals of the divergence time between different species were obtained from the TimeTree database [63].

### Identification of Hox Gene Clusters

Forty-nine unique *Hox* genes obtained from previous study were used as queries to —20 conduct BLAST search (threshold of *E* value 10^-20^) against seven species, including *P. leopoldi, C. carpio, L. oculatus, L. chalumnae, D. rerio, C. milii,* and *B. floridae,* respectively [16, 46]. The HMM profile for the homeodomain was used to identify the potential homeodomain containing genes from these genomes with HMMER 3.2.1 (http://hmmer.janelia.org/) [64], as well. All of the obtained genes were further validated using SMART database to determine whether the protein sequences contain homeodomains [65].

### Gene family expansion and contraction estimation and selective pressure analysis

Gene family expansion and contraction were analyzed by CAFE (version 3.1) using the same seven species in *Hox* gene cluster identification [66]. After filtering the genes shorter than 100 amino acids and with low sequence complexity, we collected a total of 205,728 genes from seven selected species. Orthofinder V.2.3.3 was used to classify these genes into orthologous groups based on their sequence similarity. Each orthologous group is actually a gene family. We calculated the probability of each orthologous group by 10,000 Monte Carlo random samplings and estimated the lambda value based on the maximum likelihood model, which represents the rate of expansion and contraction of each gene family. A branch with *p*-value lower than 0.05 was considered to have gene amplification and contraction over evolutionary time scales.

PAML was used to estimate the selective pressure of selected genes. Both branch and branch-site mode were applied to detected the selective pressure in *P. leopoldi* [62]. PAL2NAL was utilized for alignment nucleotide sequences based on protein alignment [67]. The likelihood ratio tests (LRTs) of M1a vs. M2a were employed to examine the selective pressure of each site among selected genes.

### Transcriptome analysis

Total RNAs from six *P. leopoldi* tissues (skin, heart, blood, muscle, liver, and brain) were prepared by using the Qiagen kit and according to the manufacturer’s instructions and RNA-seq libraries were constructed according to a standard protocol for the Illumina sequencing platform (100-bp paired-end reads). The reads were aligned onto the assembled *P. leopoldi* reference genome with STAR [68].

Other RNA-seq data of brain, heart, muscle, liver from three fishes, *L. oculatus, D. rerio, C. milii,* were retrieved from NBCI Sequence Read Archive (SRA). The raw RNA-seq data were filtered by using Trimmomatic 0.32 to generate clean reads [69]. Per-base sequence qualities of filtered fastq files were checked with FastQC (https://www.bioinformatics.babraham.ac.uk/projects/fastqc/). The Ensembl genome of each species was used as a reference genome and filtered reads were aligned onto the references by using STAR [68].

RSEM was used to quantify expressed genes into transcripts per kilobase million (TPM) value [70]. R package DEGseq was used to detect differentially expressed genes [71]. One-to-one orthologous genes between *P. leopoldi* and *D. rerio* and web-based DAVID Bioinformatics Resources were used for GO annotation [72].

### Bone-formation related gene family examination

Zebrafish’s bone morphogenetic protein (BMP) and bone morphogenetic protein receptor genes were used as query to search the BMP and BMP receptor gene families in *P. leopoldi, C. carpio, L. oculatus, L. chalumnae, C. milii,* and *B. floridae.* The other bone-formation related gene families were examined as follows. The zebrafish orthologous genes were used to annotate each gene family. The gene family with the keyword of “bone”, “calcium”, or “vitamin D” was kept as the bone-formation related candidate gene family for further examination. These bone-formation related candidate gene families were manually checked with literature evidence in order to find the target genes for knockdown experiment.

### GC gene knockdown in zebrafish

*D. rerio* (AB strain) was provided by an in-house *D. rerio* Core Facility (CAS Center for Excellence in Molecular Cell Science) and all experimental protocols were approved by the Institutional Animal Care and Use Committee. sgRNAs were designed based on the CRISPR Design website CCTop (https://crispr.cos.uni-heidelberg.de/index.html) and CHOPCHOP (http://chopchop.cbu.uib.no/). DNA templates for sgRNAs were produced by annealing and elongating a forward primer containing T7 promoter and guide sequence and a reverse primer encoding the standard chimeric sgRNA scaffold [73] (Table S12). DNA templates were purified and RNAs were *in vitro* synthesized and purified. Cas9 ribonucleoprotein (RNP) complexes were prepared with Cas9 protein and sgRNAs as previously described [74]. The RNPs were injected into one-cell stage *D. rerio* embryos. Each embryo was injected with 1 nl mix containing ~5 μM Cas9 and 1 μg/μL (31 μM) sgRNA.

*D. rerio* larvae were processed for bone staining with Alizarin Red S by using a modified protocol [75]. Briefly, 6 day-post-fertilization (dpf) larvae were fixed in 4% paraformaldehyde (PFA; w/v, pH 7.4) overnight at 4°C, washed in 1% KOH for 5 minutes, and bleached in 3%H_2_O_2_/0.5% KOH for 40 minutes. All specimens were initially stained with 0.05% Alizarin Red S in 70% ethanol overnight and soaked thoroughly with 25% glycerol/0.1%KOH, 50% glycerol/0.1%KOH and 75% glycerol/0.1%KOH sequentially. All specimens were stored in 80% glycerol/H_2_O.

## Data Availability

The raw sequencing data reported in this study have been deposited in the Genome Sequence Archive in National Genomics Data Center [76, 77], Beijing Institute of Genomics Chinese Academy of Sciences / China National Center for Bioinformation (GSA:CRA003264), and are publicly accessible at https://bigd.big.ac.cn/gsa. The whole genome sequence data and its annotation file in GFF format in this study have been deposited in the Genome Warehouse in National Genomics Data Center (GWH:GWHAOTN00000000), which are publicly accessible at https://bigd.big.ac.cn/gwh.

## CRediT authorship contribution statement

**Jingqi Zhou**: Methodology, Formal analysis, Visualization, Funding acquisition, Writing-original draft, Writing-review & editing. **Ake Liu**: Methodology, Formal analysis, Visualization, Data Curation. **Funan He**: Methodology, Formal analysis, Visualization. **Yunbing Zhang**: Formal analysis, Validation. **Libing Shen**: Conceptualization, Writing-original draft, Writing-review & editing, Supervision. **Jun Yu**: Conceptualization, Writing-review & editing, Supervision. **Xiang Zhang**: Conceptualization, Resources, Writing- review & editing, Supervision. All authors read and approved the final manuscript

## Acknowledgments

We specially thank Mr. Qiuyuan Hua of Wenzhou Hua Qiuyuan Fishery Company Limited, who provided the sample fish for this study. This study is financially supported by Shanghai Nanmulin Biotechnology Company Limited and National Science Foundation of China (Grant No. 31801049).

