## Supplemental_files for "Genome sequencing of white-blotched river stingray (*Potamotrygon leopoldi*) provides novel clues for niche-adaptation and skeleton formation": Supplemental_figure_legends.docx

**Supplemental figure 1.** The distribution of tandem repeat sequences in white-blotched river stingray genome. X axis is the divergence of stingray’s tandem repeat sequences to the TE sequences in Repbase database.

**Supplemental figure 2.** Venn diagram of the functions annotation for white-blotched river stingray’s protein coding genes using four protein databases.

**Supplemental figure 3.** The maximum likelihood tree of white-blotched river stingray and 25 selected species with statistical support values (SH-like local supports adjacent to the nodes). AMX for blind cave fish. DAR for zebrafish. GMO for Atlantic cod. GAC for three-spined stickleback. LAC for coelacanth. LOC for spotted gar. ONI for Nile tilapia. ORL for rice fish. PMA for sea lamprey. PFO for Amazon molly. TRU for Japanese puffer. TNI for green spotted puffer. XMA for platyfish. CMI for elephant shark. CSE for tongue sole. CCA for common carp. ELU for northern pike. IPU for channel catfish. NFU for turquoise killifish. OKI for coho salmon. OMY for rainbow trout. POL for Japanese flounder. PRE for guppy. SSA for Atlantic salmon. BRF for amphioxus. PLE for white-blotched river stingray. Branch length is proportional to substitution rate.

**Supplemental figure 4.** The Bayesian tree of white-blotched river stingray and 25 selected species with posterior probability support (values adjacent to the nodes). AMX for blind cave fish. DAR for zebrafish. GMO for Atlantic cod. GAC for three-spined stickleback. LAC for coelacanth. LOC for spotted gar. ONI for Nile tilapia. ORL for rice fish. PMA for sea lamprey. PFO for Amazon molly. TRU for Japanese puffer. TNI for green spotted puffer. XMA for platyfish. CMI for elephant shark. CSE for tongue sole. CCA for common carp. ELU for northern pike. IPU for channel catfish. NFU for turquoise killifish. OKI for coho salmon. OMY for rainbow trout. POL for Japanese flounder. PRE for guppy. SSA for Atlantic salmon. BRF for amphioxus. PLE for white-blotched river stingray. Branch length is proportional to substitution rate.

**Supplemental figure 5.** The species tree of 25 selected fishes and one chordate.

**Supplemental figure 5.** The MCMCtree for estimating the divergence time of white-blotched river stingray and 25 selected species. AMX for blind cave fish. DAR for zebrafish. GMO for Atlantic cod. GAC for three-spined stickleback. LAC for coelacanth. LOC for spotted gar. ONI for Nile tilapia. ORL for rice fish. PMA for sea lamprey. PFO for Amazon molly. TRU for Japanese puffer. TNI for green spotted puffer. XMA for platyfish. CMI for elephant shark. CSE for tongue sole. CCA for common carp. ELU for northern pike. IPU for channel catfish. NFU for turquoise killifish. OKI for coho salmon. OMY for rainbow trout. POL for Japanese flounder. PRE for guppy. SSA for Atlantic salmon. BRF for amphioxus. PLE for white-blotched river stingray. The numbers adjacent to the nodes are divergence time which is shown in million years (MYA).

**Supplemental figure 7.** GO enrichment analyses for expanded and contracted gene families identified in white-blotched river stingray.
