## Supplemental_files for "Genome sequencing of white-blotched river stingray (*Potamotrygon leopoldi*) provides novel clues for niche-adaptation and skeleton formation": supplemental_table_1_to_7.docx

**Table S1.** Statistics of genome sequencing data (white-blotched river stingray).

| Pair-end libraries | Insert size | Total data (G) | Read length (bp) | Sequence coverage (×) |
| --- | --- | --- | --- | --- |
| Illumina reads | 250 | 293.12 |  | 66.77 |
|  | 350 | 288.66 | 150 | 65.75 |
|  | 450 | 300.41 |  | 68.43 |
| PacBio reads | 20 Kb | 270.51 | － | 61.61 |
| 10X Genomics |  | 475.93 | 150 | 108.41 |
| Total | － | 1,628.64 | － | 370.98 |

**Table S2.** Base content statistics of white-blotched river stingray genome.

|  | Number (bp) | % of genome |
| --- | --- | --- |
| A | 1,253,750,658 | 28.77 |
| T | 1,254,374,523 | 28.79 |
| C | 914,070,220 | 20.98 |
| G | 914,572,508 | 20.99 |
| N | 20,623,492 | 0.47 |
| Total (bp) | 4,357,391,401 |  |
| GC* | 1,828,642,728 | 42.17 |

* GC content of the genome without N.

**Table S3.** Characteristic statistics of genome assembly.

| Kmer | 17 |
| --- | --- |
| Depth | 29 |
| Genome_size(M) | 4,231.24 |
| Revised Genome_size(M) | 4,209.61(4.11 gigabases) |
| Heterozygous_rate(%) | 0.79 |
| Repeat_rate(%) | 73.87 |

**Table S4.** Reads coverage statistics of white-blotched river stingray genome.

|  |  | Percentage |
| --- | --- | --- |
| Reads | Mapping rate (%) ^a^ | 98.48 |
| Genome | Average sequencing depth ^b^ | 50.91 |
|  | Coverage (%) ^c^ | 98.74 |
|  | Coverage at least 4× (%) ^d^ | 97.84 |
|  | Coverage at least 10× (%) | 95.72 |
|  | Coverage at least 20× (%) | 89.03 |

^a^ mapping rate: the percentage of reads mapping to genome.

^b^ Average sequence depth: the average sequence depth of each base mapped by reads in genome.

^c^ Coverage: the percentage of genome coverage mapped by reads.

^d^ Coverage at least NX(%): the percentage of genome coverage mapped by NX reads.

**Table S5.** Genome assembly results (white-blotched river stingray).

| Sample ID | length | | number | |
| --- | --- | --- | --- | --- |
|  | Contig**(bp) | Scaffold(bp) | Contig** | Scaffold |
| Total | 4,336,767,909 | 4,357,391,401 | 16,227 | 13,238 |
| Max | 34,999,168 | 41,069,189 | － | － |
| Number>=2000 | － | － | 15,784 | 12,796 |
| N50 | 3,937,865 | 5,675,171 | 222 | 179 |
| N60 | 2,226,993 | 3,630,306 | 367 | 275 |
| N70 | 1,057,949 | 1,818,790 | 649 | 446 |
| N80 | 359,267 | 769,167 | 1,366 | 814 |
| N90 | 113,796 | 191,036 | 3,662 | 1,982 |

**Table S6.** Statistical results of repeated sequences in white-blotched river stingray genome.

| Type | Repeat Size(bp) | % of genome |
| --- | --- | --- |
| Trf | 186,908,778 | 4.29 |
| Repeatmasker | 2,998,926,446 | 68.82 |
| Proteinmask | 62,564,836 | 1.44 |
| Total | 3,119,184,614 | 71.58 |

Note: Total repeated sequences are the results obtained by the above methods, which are the non-redundant result after removing the overlapped parts among three methods.

**Table S7.** Statistics of repeated sequence classification in white-blotched river stingray genome.

|  | Denovo + Repbase Length(bp) | % in  Genome | TE proteins Length (bp) | % in  Genome | Combined TEs Length (bp) | % in  Genome |
| --- | --- | --- | --- | --- | --- | --- |
| DNA | 19,792,076 | 0.45 | 22,913,194 | 0.53 | 42,705,270 | 0.98 |
| LINE | 2,189,913,381 | 50.26 | 61,603,347 | 1.41 | 2,251,516,728 | 51.67 |
| SINE | 721,756 | 0.02 | 0 | 0 | 721,756 | 0.02 |
| LTR | 1,064,263,616 | 24.42 | 32,380,894 | 0.74 | 1,096,644,510 | 25.17 |
| Simple repeat | 29,991,064 | 0.69 | 0 | 0 | 29,991,064 | 0.69 |
| Unknown | 4,264,845 | 0.10 | 0 | 0 | 4,264,845 | 0.10 |
| Total | 2,998,926,446 | 68.82 | 62,564,836 | 1.44 | 3,061,491,282 | 70.26 |

Note: Denovo + Repbase are the transposon elements obtained by annotating the genome using RepeatMasker software after integrating the library predicted by RepeatModeler, RepeatScout and LTR_FINDER combined with the RepBase nucleotide database using Uclust software according to the 80-80-80 principle. TE proteins are the transposon elements obtained by annotating the genome using RepeatProteinMask software based on the RepBase protein library. Combined TEs are the results of integrating the above two methods and removing redundancy. Unknown means that the repeat sequence cannot be classified by RepeatMasker.
