## Supplemental_files for "Genome sequencing of white-blotched river stingray (*Potamotrygon leopoldi*) provides novel clues for niche-adaptation and skeleton formation": supplmental_table_8.docx

**Table S8.** Basic statistical information for gene structure prediction.

|  | Gene set | Number | Average transcript length(bp) | Average CDS length (bp) | Average exons per gene | Average exon length (bp) | Average intron length (bp) |
| --- | --- | --- | --- | --- | --- | --- | --- |
| *De novo* | Augustus | 56,209 | 32,106.89 | 910.51 | 3.65 | 249.50 | 11,774.89 |
|  | GlimmerHMM | 406,617 | 9,631.17 | 376.45 | 2.68 | 140.42 | 5,506.02 |
|  | SNAP | 105,143 | 48,727.62 | 574.33 | 3.35 | 171.65 | 20,526.24 |
|  | Geneid | 66,828 | 11,027.78 | 495.14 | 3.13 | 158.19 | 4,944.57 |
|  | Genscan | 67,768 | 41,011.58 | 1,118.54 | 5.39 | 207.34 | 9,077.23 |
| Homolog | *Branchiostoma floridae* | 155,108 | 2,183.17 | 398.93 | 1.36 | 293.92 | 4,993.95 |
|  | *Callorhinchus milii* | 66,757 | 12,458.32 | 851.55 | 2.91 | 292.66 | 6,077.74 |
|  | *Danio rerio* | 47,991 | 10,643.20 | 978.27 | 2.92 | 334.45 | 5,020.84 |
|  | *Gadus morhua* | 36,147 | 15,776.59 | 861.76 | 3.46 | 249.02 | 6,061.55 |
|  | *Gasterosteus aculeatus* | 51,120 | 11,872.87 | 696.79 | 2.90 | 240.63 | 5,895.56 |
|  | *Latimeria chalumnae* | 76,153 | 7,199.62 | 924.28 | 2.21 | 417.76 | 5,175.57 |
|  | *Oreochromis niloticus* | 108,253 | 5,871.07 | 855.05 | 2.07 | 413.42 | 4,695.47 |
|  | *Oryzias latipes* | 105,441 | 6,252.47 | 838.19 | 1.98 | 423.47 | 5,528.52 |
|  | *Takifugu rubripes* | 82,069 | 9,685.26 | 870.88 | 2.36 | 369.65 | 6,500.54 |
|  | *Tetraodon nigroviridis* | 27,261 | 20,922.13 | 1,099.54 | 4.32 | 254.45 | 5,968.32 |
|  | PASA | 81,109 | 50,355.53 | 1,186.46 | 6.99 | 169.64 | 8,202.94 |
| RNAseq | Cufflinks | 70,035 | 67,076.77 | 3,854.73 | 8.19 | 470.43 | 8,788.14 |
|  | 66,210 | 21,378.76 | 814.38 | 3.55 | 229.11 | 8,049.99 |  |
| EVM | | 64,180 | 25,776.76 | 859.97 | 3.81 | 225.42 | 8,851.47 |
| Pasa-update ^a^ | | 23,240 | 57,255.91 | 1,343.22 | 7.63 | 176.02 | 8,431.71 |

^a^ It contains UTR region but no other regions.

^b^ It obtains from the result of Pasa-update after removing the variable isoforms to keep the longest transcript and removing the redundant single exon. The filtering conditions: TE overlap no less than 20%, early termination, supported only by *de novo* evidence and RPKM expression less than 1 in each tissue.
