## Supplemental_files for "Genome sequencing of white-blotched river stingray (*Potamotrygon leopoldi*) provides novel clues for niche-adaptation and skeleton formation": supplmental_table_9_to_12.docx

**Table S9.** The gene function annotation uses different methods.

|  | Number | Percent(%) |
| --- | --- | --- |
| Total | 23,240 | - |
| Swissprot | 19,440 | 83.60 |
| NR | 21,190 | 91.20 |
| KEGG | 18,385 | 79.10 |
| InterPro | 22,900 | 98.50 |
| GO | 21,040 | 90.50 |
| Pfam | 16,883 | 72.60 |
| Annotated | 23,030 | 99.10 |
| Unannotated | 210 | 0.90 |

**Table S10.** Gene structures in eleven fish species.

| Species | Number | | Average transcript length (bp) | Average CDS length (bp) | Average exons per gene | Average exon length (bp) | Average intron length (bp) |
| --- | --- | --- | --- | --- | --- | --- | --- |
| Bfl | | 28,621 | 9,236.65 | 1,392.24 | 7.03 | 198.11 | 1,301.41 |
| Cmi | | 17,847 | 24,214.70 | 1,668.28 | 9.87 | 169.06 | 2,542.48 |
| Dre | | 25,619 | 25,207.59 | 1,642.64 | 9.42 | 174.39 | 2,798.97 |
| Gac | | 20,787 | 8,451.06 | 1,548.67 | 10.4 | 148.94 | 734.44 |
| Gmo | | 20,095 | 15,245.21 | 1,459.03 | 12.72 | 114.67 | 1,175.90 |
| Lch | | 19,569 | 36,970.26 | 1,562.50 | 9.95 | 157.11 | 3,958.36 |
| Ola | | 19,699 | 12,145.58 | 1,515.82 | 10.25 | 147.82 | 1,148.61 |
| Oni | | 21,437 | 14,903.11 | 1,714.22 | 10.9 | 157.25 | 1,332.07 |
| Ple | | 23,240 | 57,255.91 | 1,343.22 | 7.63 | 176.02 | 8,431.71 |
| Tni | | 19,602 | 6,066.17 | 1,516.59 | 10.52 | 144.2 | 478.02 |
| Tru | | 18,523 | 7,492.75 | 1,693.53 | 11.1 | 152.61 | 574.33 |

Abbreviation: Bfl (*Branchiostoma floridae*), Cmi (*Callorhinchus milii*), Dre (*Danio rerio*), Gac (*Gasterosteus aculeatus*), Gmo (*Gadus morhua*), Lch (*Latimeria chalumnae*), Ola (*Oryzias latipes*), Oni (*Oreochromis niloticus*), Ple (*Potamotrygon Leopoldi*), Tni (*Tetraodon nigroviridis*), Tru (*Takifugu rubripes*).

**Table S11.** Statistics of non-coding RNAs in white-blotched river stingray genome.

|  | Type | Copy | Average length (bp) | Total length (bp) | % of genome |
| --- | --- | --- | --- | --- | --- |
| miRNA | | 1,262 | 101.18 | 127,691 | 0.002930 |
| tRNA | | 2,729 | 75.87 | 207,038 | 0.004751 |
| rRNA | rRNA | 2,405 | 176.59 | 424,687 | 0.009746 |
|  | 18S | 712 | 196.25 | 139,731 | 0.003207 |
|  | 28S | 1,653 | 170.08 | 281,136 | 0.006452 |
|  | 5.8S | 9 | 130 | 1,170 | 0.000027 |
|  | 5S | 31 | 85.48 | 2,650 | 0.000061 |
| snRNA | snRNA | 871 | 107.66 | 93,775 | 0.002152 |
|  | CD-box | 84 | 100.83 | 8,470 | 0.000194 |
|  | HACA-box | 83 | 172.92 | 14,352 | 0.000329 |
|  | splicing | 690 | 99.30 | 68,514 | 0.001572 |

**Table S12.** Primer sequences used to generate sgRNAs used in this study. sgRNA target sequences are underlined.

|  | Sequence (5'→ 3') |
| --- | --- |
| sgRNA Scaffold Primer | AAAAGCACCGACTCGGTGCCACTTTTTCAAGTTGATAACGGACTAGCCTTATTTTAACTTGCTATTTCTAGCTCTAAAAC |
| Experimental Guide  Template Primers |  |
| Control (egfp) | TAATACGACTCACTATAGGCGAGGGCGATGCCACCTAGTTTTAGAGCTAGAAATAGC |
| gc-e4 | TAATACGACTCACTATAGGCTCAATGCCTGGATGCTTGGTTTTAGAGCTAGAAATAGC |
| gc-e8 | TAATACGACTCACTATAGGTCGGTTTGGATTCATCGCAGGTTTTAGAGCTAGAAATAGC |
