## Supplementary figures and images for "Genome sequencing of white-blotched river stingray (*Potamotrygon leopoldi*) provides novel clues for niche-adaptation and skeleton formation"

### fig S5.tiff

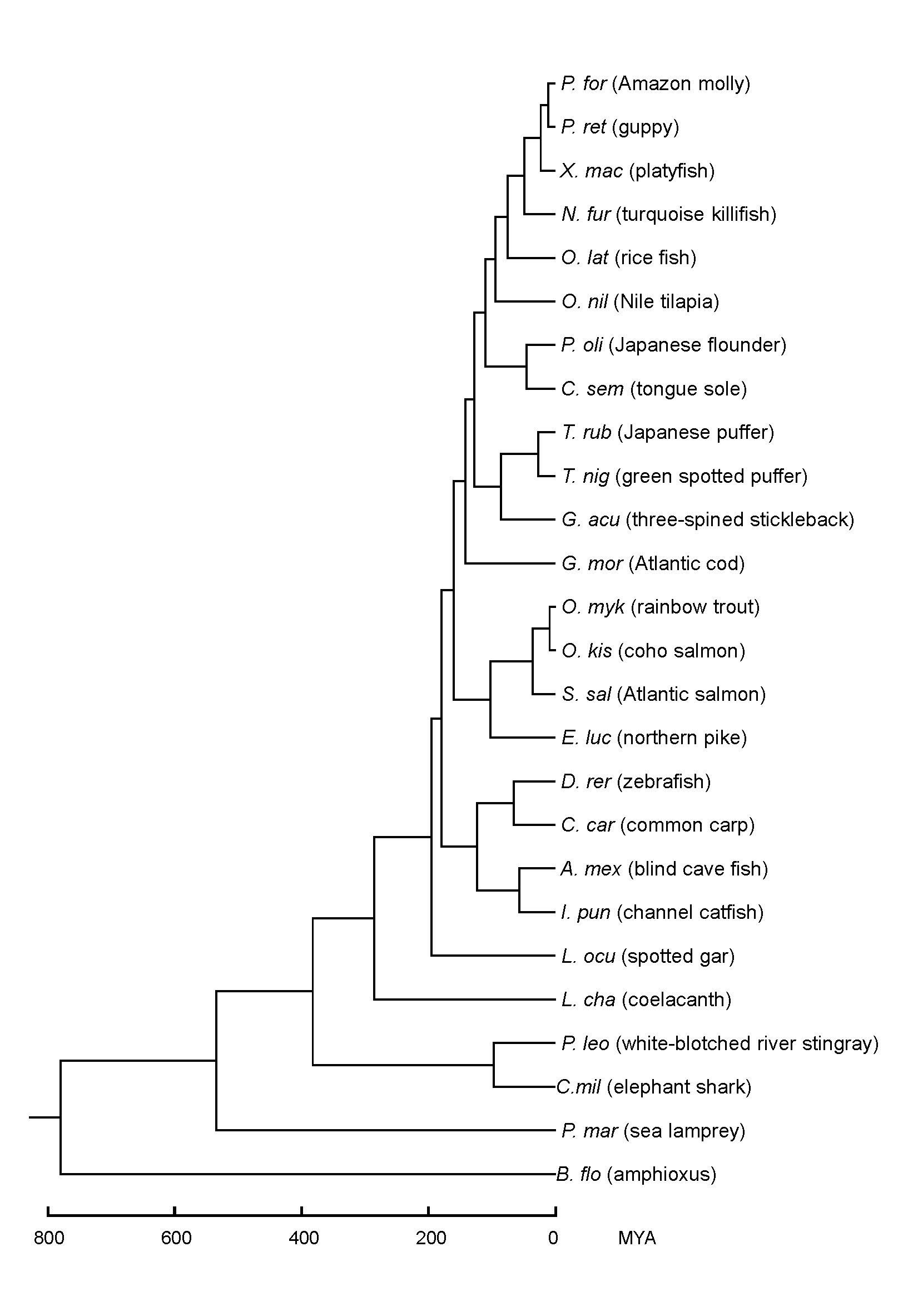

### fig S7.tiff

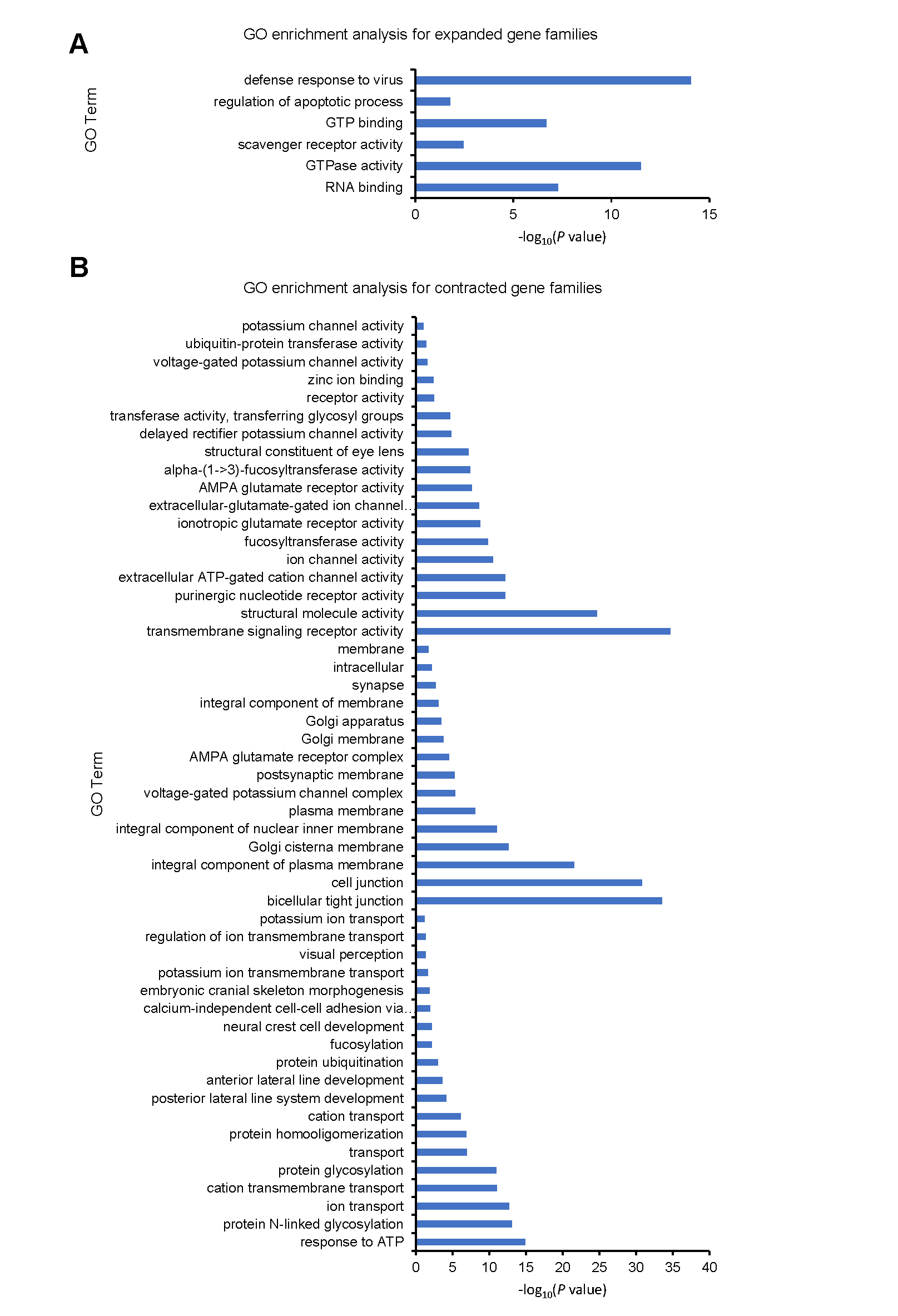

### Figure_S1.tif

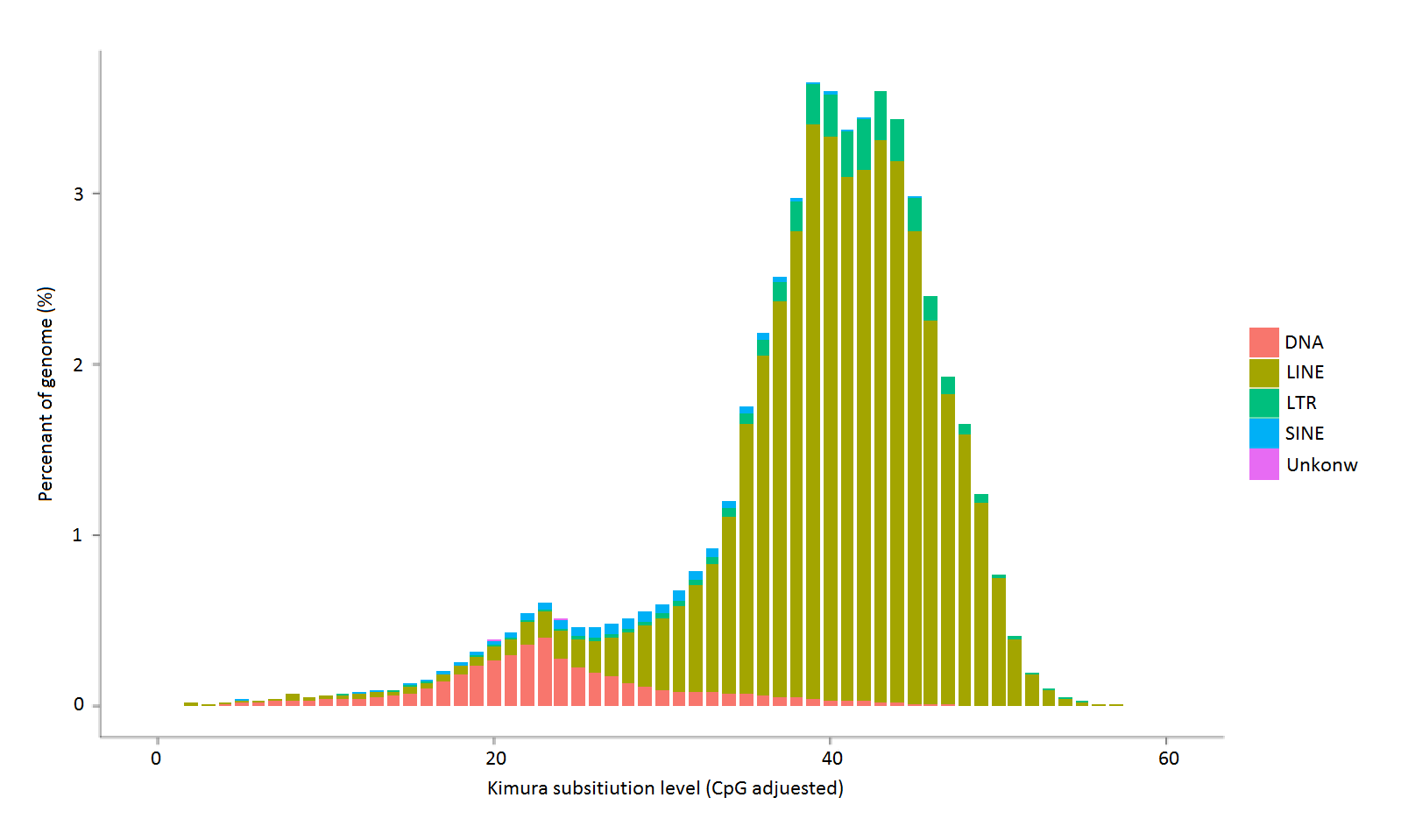

### Figure_S2.tif

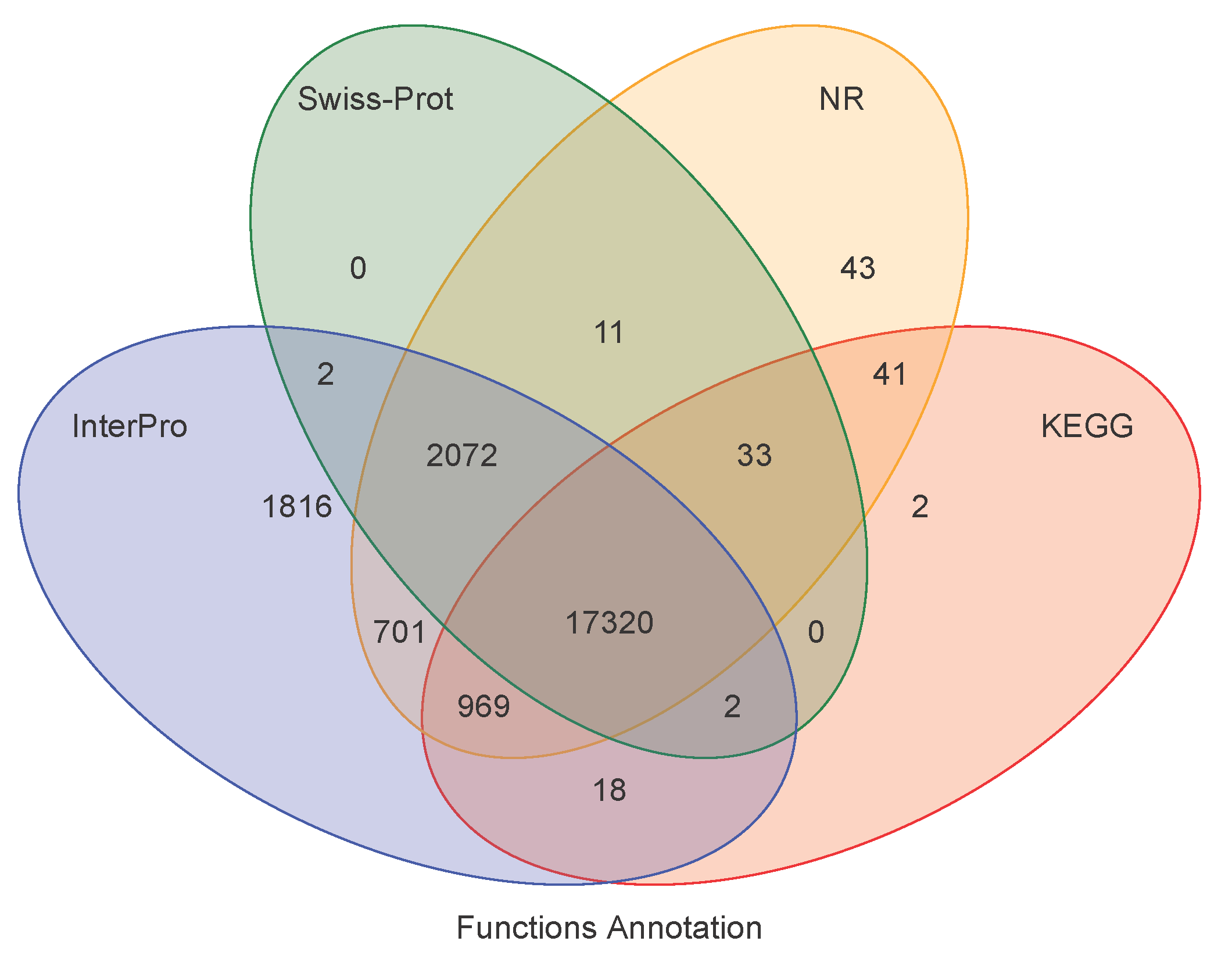

### Figure_S3.tif

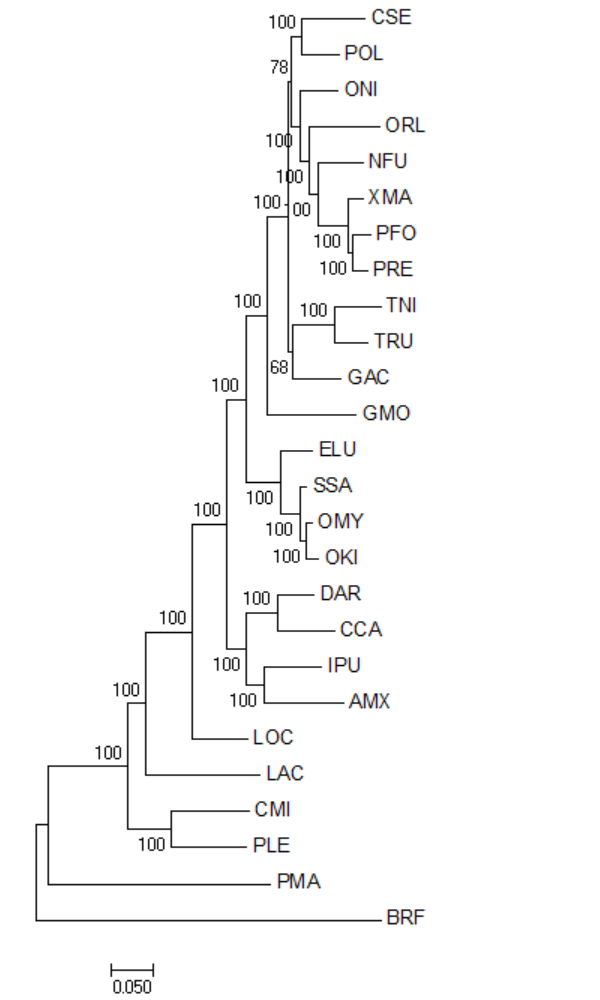

### Figure_S4.tif

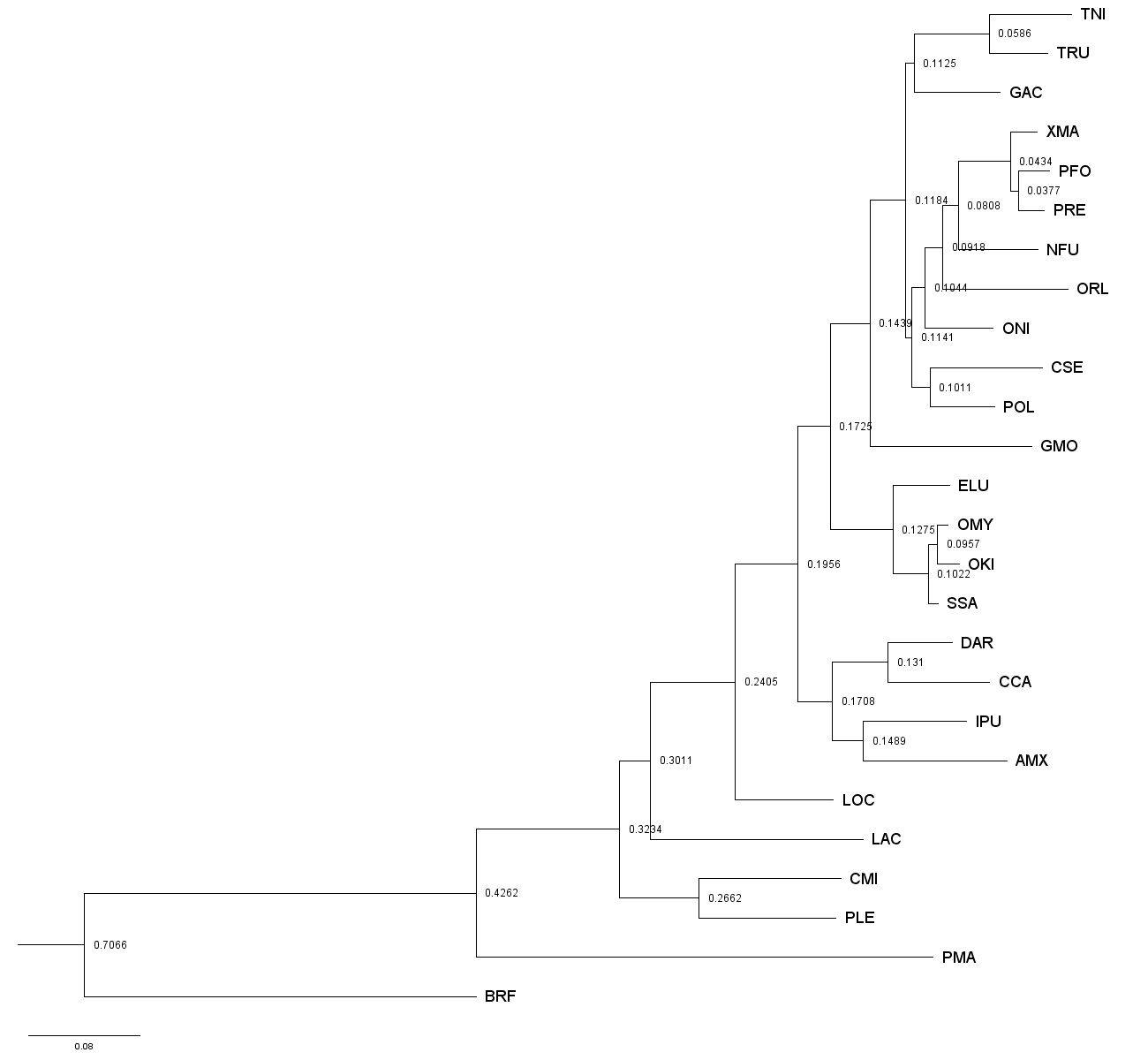

### Figure_S6.tiff

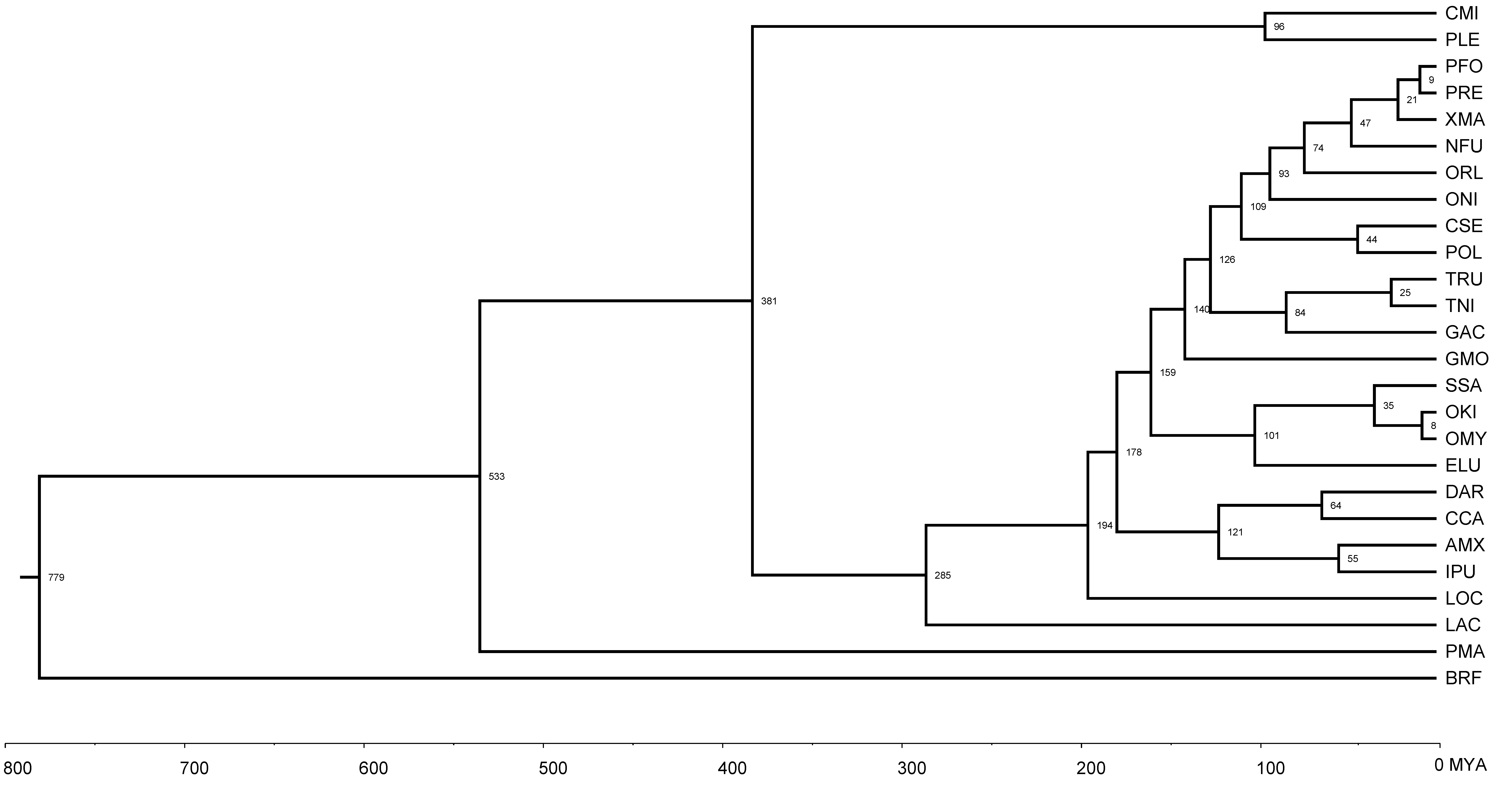
